# An adaptive autoregressive diffusion approach to design active humanized antibody and nanobody

**DOI:** 10.1101/2024.10.22.619416

**Authors:** Jian Ma, Fandi Wu, Tingyang Xu, Shaoyong Xu, Wei Liu, Divin Yan, Qifeng Bai, Jianhua Yao

**Affiliations:** School of Basic Medical Sciences, Lanzhou University, Lanzhou 730000, Gansu, P. R. China; Tencent AI Lab, Shenzhen, P. R. China; Fudan University, Shanghai, P. R. China

**Keywords:** Humanization, Antibody, Nanobody, Autogressive Diffusion Model, Deep learning

## Abstract

Humanization is a critical process for designing efficiently specific antibodies and nanobodies prior to clinical trials. Developing widely recognized deep learning techniques or frameworks for humanizing conventional antibodies and nanobodies presents a valuable yet challenging task. Inspired by the effectiveness of diffusion models across various applications, we introduce HuDiff, an adaptive diffusion approach for humanizing antibodies and nanobodies from scratch, referred to as HuDiff-Ab and HuDiff-Nb, respectively. This approach begins the humanization process exclusively with complementarity-determining region (CDR) sequences, eliminating the need for humanized templates. On public benchmark datasets, the results of HuDiff-Ab’s humanized antibodies are more similar to experimentally humanized antibodies than to those of the Sapiens humanization model. Besides, humanized nanobodies produced by HuDiff-Nb exhibit a higher humanness score and greater nativeness than those generated by the Lammanade pipeline for humanization nanobody. We apply HuDiff to humanize a mouse antibody and an alpaca nanobody, both targeting the SARS-CoV-2 RBD, and validate the binding affinity of humanized sequences through Bio-Layer Interferometry (BLI) experiments. The results show the binding affinity of the best humanized antibody is nearly equal to that of the parental mouse antibody (0.15 nM vs. 0.12 nM). Remarkably, the top-performing humanized nanobody exhibits a significantly enhanced binding affinity compared to the parental alpaca nanobody (2.52 nM vs. 5.47 nM), representing a 54% increase. These findings indicate that our approach HuDiff is highly effective in enhancing the humanness of antibodies and nanobodies while maintaining or potentially improving the binding affinity of the designed humanized sequences. The code and checkpoints of HuDiff are available at https://github.com/TencentAI4S/HuDiff.

## 1. Introduction

A conventional antibody has a characteristic Y-shaped structure with two identical pairs of protein chains, each pair consisting of a heavy chain and a light chain ^1,2^. The arms of the Y-shaped structure, known as the Fab regions, are involved in the specific recognition and binding of an antigen’s epitope. The stem of the Y-shaped structure, or the Fc region, is recognized by macrophage cells, facilitating the elimination of the antigen-antibody complex and completes the antibody’s role in the immune response^3,4^. This unique structure endows antibodies with high specificity and sufficient binding ability, resulting in an increasing number of antibody therapeutics approved by the Food and Drug Administration (FDA) for the treatment of various diseases, including cancers, autoimmune, metabolic, and infectious diseases ^5,6^. The single-domain antibody (denoted as VHH), commonly referred to as a nanobody, possesses a simpler structure that includes only a single heavy chain. Despite this difference, nanobodies can perform functions similar to conventional antibodies ^7^. Notably, The complementarity-determining region (CDR), particularly CDR3, of a VHH is often longer than that of a conventional antibody heavy chain (VH) ^8^. Nanobodies are attracting growing interest due to properties such as small size, high solubility, thermal stability, and effective tissue penetration in vivo ^9–11^. These attributes give nanobodies distinct advantages, such as applications in brain therapy (crossing the blood-brain barrier), tumor imaging, and nanobody-drug conjugates ^12,13^. Nanobodies are also being explored in precision medicine tailored to individual patients ^14–16^. However, in clinical practice, it has been observed that if a monoclonal antibody is not recognized by the human immune system, a human anti-mouse antibody (HAMA) response can occur ^17,18^. This response can diminish the effectiveness of the murine antibody. Consequently, humanizing antibodies derived from mice or nanobodies from camels is a critical step before they can advance to clinical trials ^19,20,5^.

Humanization of mouse-derived antibodies and camel-derived nanobodies remains an important and challenging area of research. The primary goal of humanization is to reduce or entirely eliminate the immune reactions that these therapeutic agents may provoke. It is hoped that humanized antibodies will retain a strong affinity for their target antigens, without experiencing a notable reduction in binding efficacy compared to their original versions ^21,22^. To this end, various traditional methods have been developed, most of which focus on the gene type. One such approach is the production of chimeric antibodies ^23,24^, where the variable regions are derived from mouse antibodies, and the constant regions come from human antibodies. While the chimeric antibody retains its antigen-binding affinity, the mouse-derived variable region may still provoke an immune response. Another approach is CDR grafting, which involves transferring the complementarity-determining regions (CDRs) from mouse antibodies or nanobodies into the variable framework of a human antibody template. This substantially reduces the likelihood of the murine antibody or camel nanobody triggering human immune responses ^25–27,21,7^. The process requires selecting a framework region (FR) template from a library of human sequences that closely matches the mouse FR sequence. The goal is to limit alterations to the FR sequence to preserve the integrity of CDRs. In addition, researchers have introduced a technique known as backmutation, which posits that certain residues, particularly Vernier residues, at specific positions are crucial for maintaining the CDR structure ^28–30^. Following CDR grafting, backmutation is performed to restore any critical residues that support the CDR’s conformation. However, since CDRs originate from mice, there remains a risk of triggering anti-idiotypic reactions in patients. To further diminish this risk, the grafting of specificity-determining residues (SDRs) ^31–34^ has been proposed. This method retains only the residues in the CDRs that directly interact with the antigen, while replacing the remaining residues with those from human antibodies. Its success relies heavily on the structural analysis of the mouse antibody-antigen complex to identify which CDR amino acids should be preserved. Consequently, the applicability of SDR grafting is limited by the constraints of structural analysis.

With advances in research, a wide range of deep learning models, statistical methods, and structural strategies have been developed for antibody and nanobody humanization. For antibodies, the deep learning model Humab and the statistical method MG both employ a humanness score threshold to guide the antibody humanization process ^35,36^. Another model, Sapiens ^37^, like our approach, is a sequence-based method that employs two distinct transformer-based models—one for the heavy chain and one for the light chain—trained on human antibody sequences to directly humanize the framework region (FR) of an antibody without relying on a scoring system. The CUMAb method, introduced to account for antibody structure ^38^, creates a library of three-dimensional (3D) antibody framework structures, integrates the mouse CDRs to form the complete 3D antibody, and selects the structure with the lowest energy as the humanized antibody. For nanobodies, a pipeline called Llamanade ^39^ has been developed that uses statistical analysis to identify residue frequency differences between VH (heavy chain of antibody) and VHH while incorporating structural information for humanization. Similarly, AbNativ ^40^ introduced a pipeline that employs two models: the VH-humanness model to assess the impact of residue changes on the humanness score, and the VHH-nativeness model to prevent excessive modifications that could compromise the nativeness score of the nanobody. Besides, a variety of scoring systems have been established to assess the degree of humanization in antibodies and nanobodies, such as T20, MG score, H score, and G score, which are derived from extensive analysis of variable regions in human antibody datasets. These scoring systems utilize diverse approaches, including heuristic equations, statistical computations, and deep learning algorithms ^41,36,42,43^, which excel at identifying sequences that either closely mirror or deviate from human sequences. The Oasis database ^37^, created from 9-mer peptides mined from human antibody datasets, aids in the humanization assessment of engineered antibodies. Furthermore, deep learning approaches such as ABLSTM^44^ and AbNativ ^40^, which are trained on datasets of human antibodies, have demonstrated impressive accuracy on test sets comprising antibodies from various sources.

Diffusion models ^45,46^ have recently been applied across a wide range of tasks ^47^, achieving notable successes, particularly in biological research. For example, in protein structure prediction, AlphaFold3^48^, a leading model, integrates a diffusion component to enhance the accuracy of atomic position predictions across all molecules. To further investigate the relationship between sequence and structure in protein design, ProteinGenerator ^49^ and RFdiffusion ^50^, both developed based on RoseTTAfold ^51^, have been introduced. These two models facilitate the novel creation of protein structures, showcasing the capacity of diffusion models to comprehend complex biomolecular architectures. In contrast, given the existing vast sequence space of evolutionary-scale sequence data, Evodiff ^52^ and DPLM ^53^ perform protein design at the sequence level through a diffusion process. Building on these advances and the extensive data available for antibody and nanobody sequences, it can be inferred that diffusion models may be effectively utilized for the design of humanized antibodies and nanobodies.

In this study, we introduce an adaptive and efficient diffusion approach for the humanization of antibodies and nanobodies in sequence space, which only requires CDR sequences, eliminates the need for a pre-existing humanized framework template, and differs from methods relying on mutation pipelines to achieve predefined values. The proposed method comprises a pre-training step utilizing human sequences and a fine-tuning step for humanizing target sequences, termed HuDiff, which facilitates the humanization of antibodies (denoted as HuDiff-Ab) and nanobodies (denoted as HuDiff-Nb). Notably, unlike other methods that focus on humanizing individual antibody chains, HuDiff-Ab humanizes both the heavy and light chains simultaneously, potentially taking inter-chain information into account during the process. We evaluate HuDiff-Ab and HuDiff-Nb on various public datasets. Regardless of whether the test sets involve antibodies or nanobodies, the results demonstrate improved humanness in the humanized antibodies and nanobodies produced in silico. Using HuDiff-Ab and HuDiff-Nb, we humanize the murine antibody 2B04 and the alpaca nanobody 3-2A2-4^54^, both of which target the SARS-CoV-2 Spike RBD protein ^55^, resulting in five humanized antibodies and five humanized nanobodies. In Bio-Layer Interferometry (BLI) experiments, we observe that the affinity of the best humanized antibody is nearly identical to that of the murine antibody (0.15 nM vs. 0.12 nM), while other humanized antibodies retain their binding activity. Similarly, all humanized nanobodies preserve their binding affinity, with the best humanized nanobody showing even stronger affinity than the original alpaca nanobody (2.52 nM vs. 5.47 nM). Collectively, these findings demonstrate that HuDiff enhances the humanness of antibodies and nanobodies while preserving or even improving their binding affinity, suggesting that HuDiff could potentially serve as a valuable method for diverse humanization tasks.

## 2. Results

In the following sections, we first describe the application of HuDiff for humanizing antibodies and nanobodies (see section 2.1), then evaluate the performance of HuDiff-Ab through in silico analyses and validate the humanized antibodies with wet lab experiments (see section 2.2). Finally, we demonstrate the advantages of HuDiff-Nb, which is similarly validated through wet lab experiments for humanized nanobodies (see section 2.3). In both training processes, the complementarity-determining region (CDR) residues remain unchanged throughout the forward process, from timestep 0 to T, regardless of whether the sequences originate from antibodies or nanobodies (depicted in Figure 1a). During this process, the distribution of framework region (FR) sequences evolves into a uniform distribution (absorbing state). By timestep T, the sequence retains only the CDR residues, with no remaining information about the FR residues. After training, our models act as denoisers, retrieving the CDR sequences and reconstructing the FR sequences from the uniform distribution, thereby completing the humanization process.

**Figure 1.**
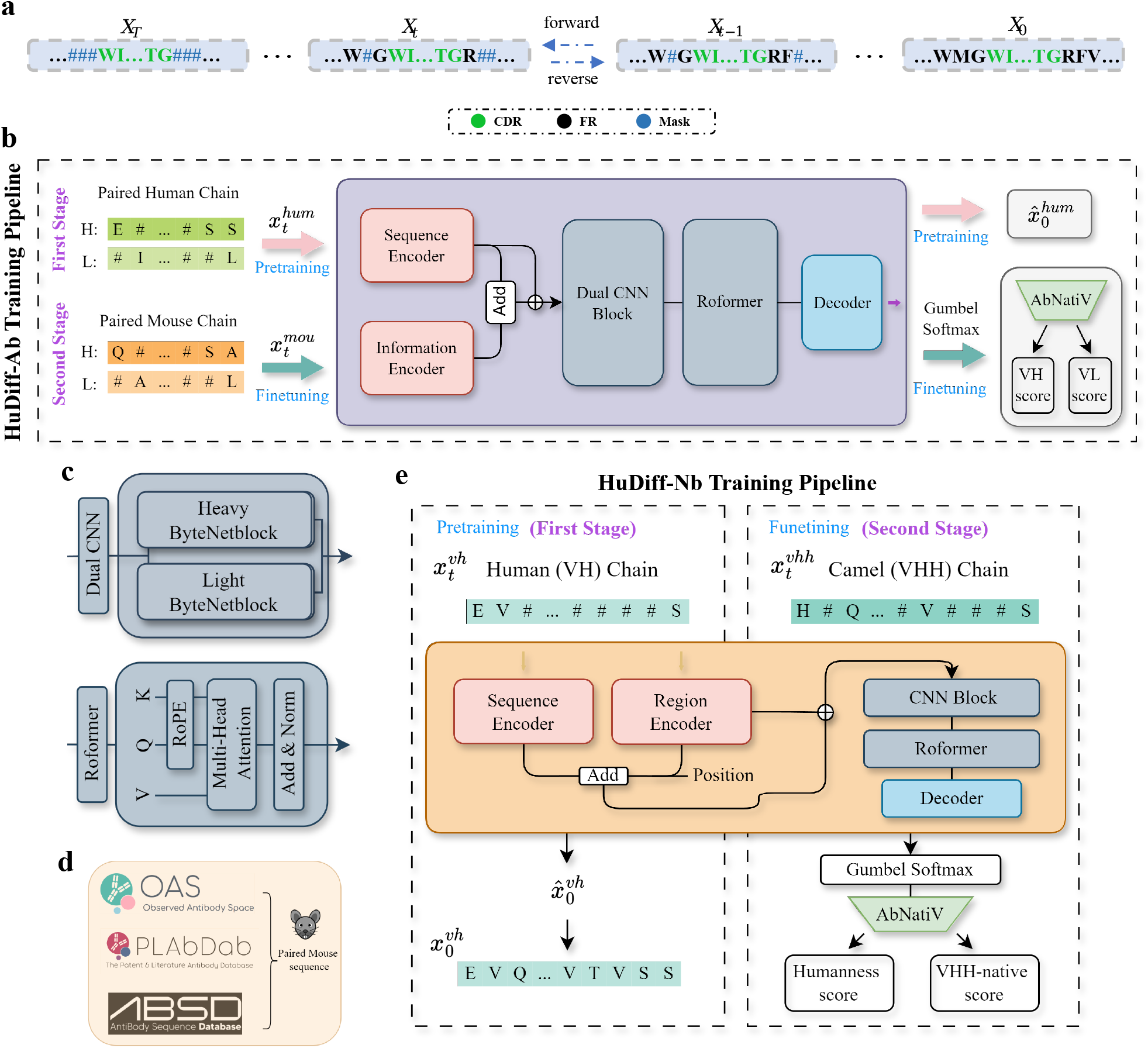
Diffusion process and model architecture. **a**, The diffusion process in models. Green text indicates the CDR residues, which remain unchanged throughout the process. Black text represents the FR residues that stay consistent at each timestep *t*. Blue text signifies the masked (or diffused) state of the FR residues at each timestep *t*. In the forward process, all FR residues become diffused, except for the CDR residues. During reverse, denoisers restore the FR residues from their abstract state. **b**, The architecture of HuDiff-Ab training pipeline. In the pre-training stage, the model takes paired human heavy and light chain sequences at various diffusion time steps *t*, with the output 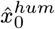 approximating the human antibody at time step 0. In the fine-tuning stage, the input consists of paired mouse antibody sequences at different diffusion time steps. The output is transformed into a one-hot vector using Gumbel softmax trick and evaluated by the AbNatiV model, producing VH and VL scores. **c**, The architecture of the Dual CNN block and the Roformer Block. **d**, The paired mouse antibody sequences used in this study are sourced from the OAS, PLAbDab, and ABSD databases, while all other training sequences are sourced from the OAS. **e**, The architecture of HuDiff-Nb training pipeline. In the pre-training phase, the model takes diffused human heavy chain sequences as input and generates denoised sequences. During the fine-tuning phase, the model uses camel nanobody sequences to produce humanized nanobodies. This output is transformed into a one-hot vector via Gumbel softmax trick, and is subsequently evaluated by AbNatiV model to predict the Humanness score (VH score) and VHH-native score, which guide the training objective during this phase.

### 2.1. HuDiff-Ab and HuDiff-Nb

#### Architecture and training pipeline of HuDiff-Ab

As shown in Figure 1b, our HuDiff-Ab training pipeline comprises two stages. In the first stage, the input to HuDiff-Ab is the masked state (diffused state) 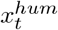 of the paired human antibody, where *t* is determined by the framework length of the paired sequence. The output is 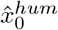, which represents the sequence predicted by the model for the masked positions at various diffusion time steps. This pre-training stage enables the model to discern the intricate relationships between known sequences in different diffusion states and the target sequences requiring reconstruction, aligning closely with our objective to design the framework region sequence. In the second stage, we use the paired mouse sequences to fine-tune HuDiff-Ab, allowing the model to further learn the humanization changes from mouse to human. Thus, the input is the diffused state 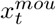, and the output logits of sequences are converted into one-hot vectors using Gumbel Softmax. These vectors are subsequently evaluated by the VH and VL models from AbNativ, which generate humanness scores for the residues in the output sequences. Our training objective is to maximize these scores (see Methods). Additionally, using Gumbel Softmax allows for gradient backpropagation during the fine-tuning process.

The architecture of HuDiff-Ab consists of three primary components, as illustrated in Figure 1b. The first component is the encoders, which encompass both a sequence encoder (see Supplementary Figure S1a)and an additional information encoder. Given that our residue inputs are numerical tokens, the sequence encoder translates these tokens into hidden features. However, the sequence encoder alone maps only the features of individual residues. To address this limitation, we incorporate the information encoder, which comprises a region encoder, a position encoder, and a chain type encoder (see Supplementary Figure S1b). The region encoder provides HuDiff-Ab with information about specific CDR or FR regions within the paired sequence. The position encoder introduces distance information to facilitate the model’s understanding of the relationships between residues and their positions. The chain type encoder determines whether the paired sequence is of the H-K or H-L type, with H denoting the heavy chain, thereby recognizing the subtle distinction between Lambda (L) and Kappa (K) light chains. Collectively, these two encoders deliver a comprehensive feature representation of the paired chain to the model. The second component is the hidden feature study blocks, which enable HuDiff-Ab to discern more complex relationships. We employ a Dual CNN block (see Figure 1b) to capture locally significant information within each individual chain. This block consists of N heavy ByteNetBlocks and N light ByteNetBlocks (see Figure 1c). When the encoder features enter the Dual CNN block, the paired sequence is divided into heavy and light chains based on their lengths. The features of each chain are independently mapped using these parallel ByteNetBlocks, and the results are concatenated along the sequence length dimension. To further model the relationships between the paired sequences, we utilize a Roformer Block (illustrated in Figure 1b and c) to learn the intricate relationships among different segments of the entire paired sequences, enabling HuDiff-Ab to accurately identify the interconnections among the six CDRs and eight FR sequences. The third component is the decoder, which employs Multilayer Perceptron (MLP) to translate the high-order features into final predictions of residues.

Unlike other models or methods, HuDiff-Ab is trained using paired sequences, as it is believed that simultaneous modeling of paired sequences is essential for humanizing a mouse antibody, given that an antibody consists of both heavy and light chains. This perspective is supported by experiments demonstrating that paired antibody data can enhance the performance of antibody language models ^56^. In our pre-training stage, we exclusively utilize preprocessed human paired sequences from the OAS database ^57^ (see Methods). During the fine-tuning phase for mouse antibodies, as shown in Figure 1d, we collect data from the OAS, PLAbDab ^58^, and ABSD database. Furthermore, all training stages involve aligning the paired sequences using the IMGT numbering scheme ^59^.

#### Architecture and training pipeline of HuDiff-Nb

Before training the HuDiff-Nb, we analyze the differences between human heavy chains (VH) and camelid heavy chain single domains (VHH). We first examine the variance in distribution patterns (see Supplementary Figure S2) between VH CDR sequences and VHH CDR sequences, revealing significant differences, particularly in the CDR3 region. We assess the length frequency (see Supplementary Figures S3 and S4), showing notable differences in the most common CDR3 and CDR2 lengths between VH and VHH sequences. These findings indicate that the length and residues of CDRs differ between VH and VHH sequences, representing an out-of-distribution problem. Fortunately, our approach addresses this issue by incorporating CDRs information from VHH sequences in the second fine-tuning stage. Our in silico humanness experiments (see section 2.3) demonstrate that the fine-tuned HuDiff-Nb outperforms its pretrained version.

The architecture of HuDiff-Nb is similar to HuDiff-Ab, which can be divded as three componets of encoders, hidden blocks and a decoder (see Figure 1e). Due to the absence of the light chain for VHH, the Information encoder is modified to region encoder, in contrast to HuDiff-Ab. Additionally, in the hidden block part, the HuDiff-Nb contains a CNN Block instead of the Dual CNN block used in HuDiff-Ab. The training process for HuDiff-Nb follows a pipeline similar to that of HuDiff-Ab and consists of two main stages (see Figure 1e). During the pre-training stage, the model is designed to reconstruct the VH sequences. This involves training with corrupted VH sequences at various timesteps and predicting the 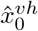 which is closer to the true 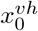. In this stage, the model captures the relationships between the CDR and FR of human sequences. In the fine-tuning stage, the model works with corrupted VHH sequences at different timesteps. For the output, similar to the HuDiff-Ab, we incorporate the VH and VHH models of AbNativ (see Methods). The VH model assesses the humanness score of each residue in a sequence, while the VHH model evaluates the nativeness score that how well the residues fit the VHH context. Unlike HuDiff-Ab, which aims to maximize both scores, the HuDiff-Nb focuses solely on maximizing the humanness score while maintaining the VHH nativeness score relatively stable (see Methods). Furthermore, we continue to employ Gumbel-Softmax trick to ensure that gradients can be backpropagated, similar to the process in HuDiff-Ab. The pre-training antibody heavy chain dataset and the fine-tuning nanobody dataset are both sourced from the OAS database. All sequences are aligned using the IMGT numbering scheme, similar to the approach used for HuDiff-Ab (see Methods).

### 2.2. Evaluation of humanness performance and experimental validation of HuDiff-Ab

#### Humanization process and performance of HuDiff-Ab

We employ various public datasets to evaluate the efficacy of HuDiff-Ab in antibody humanization. Here, we primarily present the model’s performance on the HuAb348 and Humab25 humanization datasets. Figure 2a illustrates the humanization process of HuDiff-Ab, which requires only the CDRs from both the heavy and light chains as inputs. HuDiff-Ab designs the FR sequence through reconstructing the masked residues by T iterative time steps. For benchmarking, we select the widely recognized Sapiens model and use its evaluation metrics (OASis and T20 scores) to assess improvements in humanness. Additionally, we introduce a new metric called Germline Identity, which measures the similarity between the framework region (FR) sequences of the humanized antibodies and those of the closest human germline. Table 1 presents the humanness performance of humanized antibodies generated using traditional grafting method, the Sapiens model, and the HuDiff-Ab across test datasets. The traditional grafting method, which directly substitutes mouse framework regions (FR) with the closest human germline FR, shows higher humanness scores. However, this method often results in a significant reduction in affinity, highlighting the trade-off between humanness and affinity retention. Thus, higher humanness does not always correlate with improved outcomes. Based on experimental results, achieving humanness scores that are balanced may lead to better overall performance. The humanness improvements achieved by the HuDiff-Ab and Sapiens are notably close to those of experimentally humanized antibodies. The term HuDiff-Ab of Tabel 1 refers to the process of humanizing mouse antibodies once, using a shuffled sampling order. Our findings indicate that HuDiff-Ab surpasses ‘Sapiens1’ in both OASis and T20 scores on the Humab25 dataset, demonstrating its effectiveness in designing human-like antibodies from scratch. Although HuDiff-Ab does not achieve OASis scores as close to the experimental data as Sapiens model on the HuAb348 dataset, its T20 score is more closely aligned with the experimental results. Unlike Sapiens, which can further humanize its own previous results, HuDiff-Ab is for one-stop generation. Therefore, we integrate the evaluation metrics to select desired generated results from multiple runs. HuDiff-Ab_oasis_ iterates the humanization process ten times, selecting the antibody with the highest OASis score. Similarly, HuDiff-Ab_T20_ chooses the top antibody out of ten based on the T20 score. In contrast, HuDiff-Ab_simi_ focuses on selecting antibodies that most closely resemble the original mouse antibody. HuDiff-Ab_oasis_ and HuDiff-Ab_T20_ demonstrate greater improvements in human-likeness on the OASis and T20 metrics than any iterative steps of the Sapiens. HuDiff-Ab_simi_ selects antibodies most similar to mouse antibodies, achieves high improvements on both OASis and T20 while maintaining a high Germline identity. This highlights the effectiveness of our approach in humanizing mouse antibodies. Notably, even without using FR sequence information, HuDiff-Ab achieves a higher heavy chain identity than all iterative steps of Sapiens, with its humanness improvement more closely aligning with experimental values. Overall, the superior performance of our methods on both T20 and OASis scales highlights their potential for generating antibody sequences that closely resemble human antibodies.

**Table 1.** Comparison of humanness performance across different methods in various public test datasets. The humanness improvement, such as an increase of 29.91%, refers to the absolute change in the humanness score, not a relative change. The Oasis value represents the absolute change in OASis medium identity between humanized antibodies and their corresponding parental antibodies. T20 score represents the absolute change between humanized antibodies and their parental counterparts. Germline identity refers to the sequence similarity between the humanized antibodies and the most human germline sequence, as determined by ANARCI. Bold values indicate that, for this metric, the performance of the in silico humanized sequences is closest to that of the experimentally validated sequences.

| Data | Method | Humanness improvement |  | Germline identity |  |
| --- | --- | --- | --- | --- | --- |
|  |  | Oasis | T20 | FR |  |
|  |  |  |  | H | L |
| HuAb348 | Experiment | 29.91% | 11.30% | 89.51% | 91.72% |
|  | Sapiens1 | 29.67% | 9.85% | 86.06% | 89.43% |
|  | Sapiens2 | 33.20% | 11.84% | 88.86% | 91.57% |
|  | Sapiens3 | 34.00% | 12.19% | 89.74% | 92.08% |
|  | straight graft | 35.13% | 14.55% | 94.71% | 96.37% |
|  | vernier graft | 30.35% | 12.15% | 90.79% | 94.49% |
|  | HuDiff-Ab | **31.93%** | **11.32%** | 91.44% | 87.43% |
|  | HuDiff-Ab<sub>oasis</sub> | 34.41% | 11.62% | 91.80% | 88.62% |
|  | HuDiff-Ab<sub>T20</sub> | 33.36% | 12.47% | 92.25% | 89.78% |
|  | HuDiff-Ab<sub>simi</sub> | 32.48% | 11.57% | 91.46% | 88.32% |
| Humab25 | Experiment | 33.82% | 13.1% | 89.36% | 94.47% |
|  | Sapiens1 | 29.70% | 10.20% | 84.61% | 89.75% |
|  | Sapiens2 | 33.30% | 11.80% | 87.30% | 91.46% |
|  | Sapiens3 | 34.00% | 12.40% | 88.48% | 91.87% |
|  | straight graft | 36.17% | 14.56% | 94.59% | 96.27% |
|  | vernier graft | 32.00% | 12.12% | 91.00% | 94.88% |
|  | HuDiff-Ab | **33.07%** | 11.46% | 90.32% | 87.96% |
|  | HuDiff-Ab<sub>oasis</sub> | 35.25% | 11.31% | 91.16% | 89.13% |
|  | HuDiff-Ab<sub>T20</sub> | 34.05% | **12.89%** | 91.25% | 90.16% |
|  | HuDiff-Ab<sub>simi</sub> | 33.04% | 11.10% | 90.59% | 88.63% |

**Figure 2.**
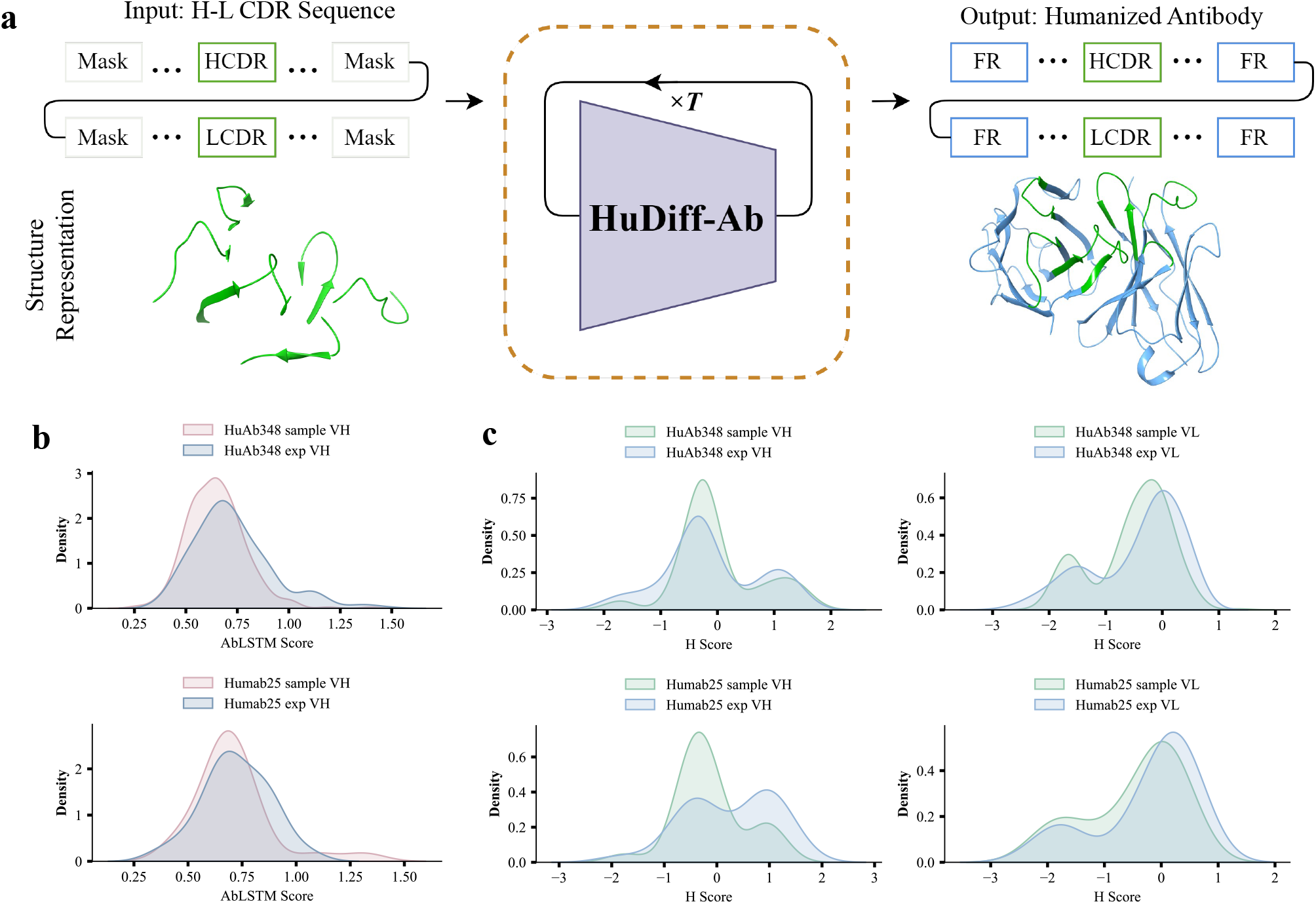
Humanization process and performance of HuDiff-Ab model. **a**, This diagram outlines the humanization process of HuDiff-Ab model. The model initially takes in six CDR sequences (green blocks) from paired H-L sequences along with the masked states of the FR sequences (white blocks). After T iterations, corresponding to the length of the masked (diffused) sequence, the humanized FR sequences (blue blocks) are combined to produce a humanized nanobody. This process mirrors the structural representation, starting with the CDRs and culminating in the completion of the FR structure. **b**, Distribution comparison of AbLSTM scores between our humanized sequences and experimentally humanized sequences across different test datasets. ‘sample’ refers to sequences humanized by our model, while ‘exp’ denotes experimentally humanized sequences. **c**, Comparison of the distribution of H scores between experimentally humanized heavy and light chains and those generated by our model across various test datasets.

Following the methodology outlined by BioPhi ^37^, we evaluate the similarity between our humanized antibodies and their experimental counterparts, focusing on two key metrics: preservation and mutation precision, as detailed in the Methods section. In summary, preservation measures the extent to which residues from the original mouse sequences are retained in the humanized antibodies, while mutation precision assesses the consistency of mutated residues between the humanized antibodies and experimental results. For a fair comparison, we select only Sapiens1 and the direct humanization results of HuDiff-Ab, neither of which involve multiple humanization processes. Table 2 presents the results for two distinct public test sets. ‘Total’ refers to all residues in the sequence, while ‘Vernier’ denotes the key residues defined by the Kabat numbering scheme ^60^. In terms of preservation, achieving closer alignment with experimental results is preferable. As expected, our findings indicate that our method exhibits a closer alignment with experimental results regarding total preservation compared to Sapiens1. However, in terms of vernier preservation, our method performs slightly worse than the Sapiens model. Similarly, our method exhibits lower mutation accuracy compared to Sapiens. These decreases are anticipated due to the nature of our approach, which does not incorporate the original framework region (FR) sequences, whereas the Sapiens does include them. As a result of these differences, HuDiff-Ab may generate a more diverse array of humanized antibody sequences, which could be advantageous in applications where diversity is desired.

**Table 2.** Sequence preservation and similarity between humanized and experimental antibodies. The table reports sequence preservation as the percentage of identity between the parental and humanized sequences, both ‘Total’ and ‘Vernier’. ‘Total’ refers to all the residues in a sequence, while ‘Vernier’ denotes the key residues defined by the Kabat numbering scheme. Mutation Precision measures the number of consistent mutations between the predicted and experimental humanized sequences, also categorized as Total and Vernier. Germline Identity quantifies the similarity between the humanized framework region sequences and the closest human germline sequences. Bold values indicate that, for this metric, the performance of the in silico humanized sequences is closest to that of the experimentally validated sequences.

| Data | Method | Preservation |  |  |  | Mutation precision |  |
| --- | --- | --- | --- | --- | --- | --- | --- |
|  |  | Total |  | Vernier |  | Total | Vernier |
|  |  | H | L | H | L |  |  |
| HuAb348 | Experiment | 82.86% | 84.50% | 83.53% | 93.00% | - | - |
|  | Sapiens1 | 88.85% | 88.43% | 86.04% | 91.41% | 69.70% | 46.53% |
|  | HuDiff-Ab | **83.28%** | **82.32%** | 77.60% | 89.46% | 57.33% | 35.03% |
| Humab25 | Experiment | 78.34% | 82.35% | 79.25% | 93.14% | - | - |
|  | Sapiens1 | 88.54% | 88.33% | 86.00% | 94.00% | 77.00% | 49.00% |
|  | HuDiff-Ab | **82.13%** | **82.49%** | **79.16%** | 91.14% | 60.85% | 38.68% |

To enhance the evaluation of antibody humanization approach, AbLSTM score and H score are introduced. These evaluations focus on the humanized antibodies derived from the HuDiff-Ab results. Since AbLSTM score assesses only heavy chains, our analysis is limited to this component. Figure 2b shows that the distribution of AbLSTM scores for our humanized heavy chains closely matches those of the experimental counterparts, with similar results observed across both the HuAb348 and Humab25 datasets. In Figure 2c, we present the H score distributions for both the experimental and in silico humanized antibody pairs. For the HuAb348 dataset, the distributions of both heavy and light chains for our humanized antibodies closely resemble those of the experimental antibodies. In the Humab25 dataset, the light chain distribution of our humanized antibodies is comparable to the experimental data, while the heavy chain distribution, although not closely matched, still demonstrates significant overlap.

#### Humanization of antibody 2B04 targeting SARS-CoV-2 RBD and experimental validation

The murine antibody, named 2B04, shows strong neutralization against SARS-CoV-2, and the complex crystal structure of Fab 2B04 bound to SARS-CoV-2 receptor-binding domain (RBD) is experimentally solved ^61^. To simulate the antibody humanization process and evaluate the effectiveness of HuDiff-Ab, we select this murine antibody. HuDiff-Ab generates numerous humanized variants, from which five are ultimately chosen for production and in vitro experiments. As shown in Figure 3a and 3b, the humanness of the five selected humanized antibodies is evaluated. Figure 3a shows Oasis Medium Score for all humanized antibodies alongside 2B04, demonstrating that the scores of our humanized antibodies exceed that of 2B04. To further confirm humanness, we use VH and VLambda models of AbNativ to assess the VH and VL scores for the heavy and light chains, respectively. As illustrated in Figure 3b, all humanized antibodies scored above the threshold value of 0.8, indicating greater humanness than 2B04. Additionally, the human germline identity for both the heavy and light chains is calculated, revealing higher germline identity in all humanized antibodies compared to 2B04 (see Figure 3c). These results suggest that our humanized antibodies are indeed human-like and may elicit a lower immune response..

**Figure 3.**
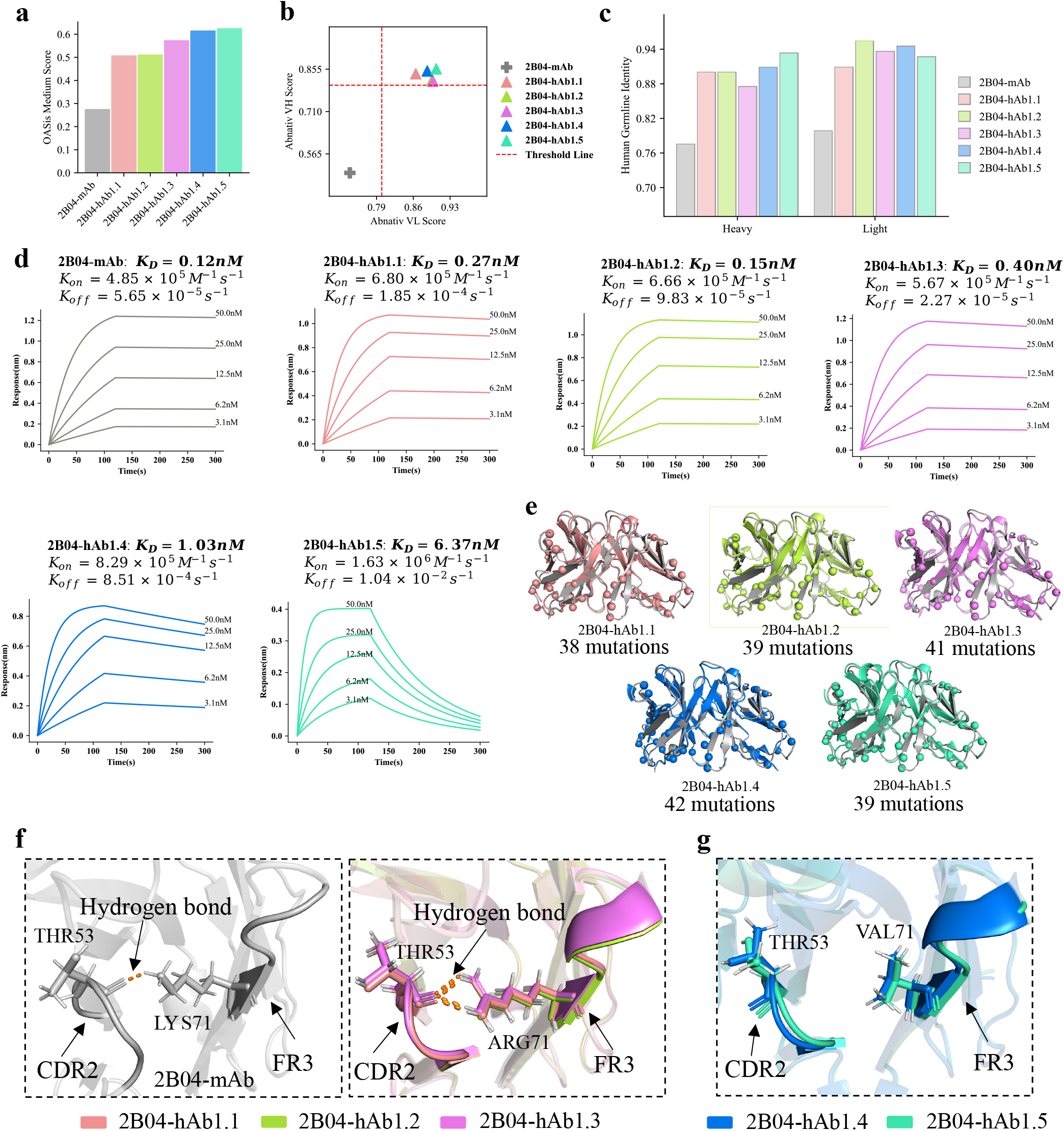
Humanness and binding affinity of humanized antibodies. **a**, Comparison of OASis Medium scores of our five humanized antibodies with those of the parental mouse antibody (grey bar). Higher values indicate greater humanness according to this metric. **b**, Displaying the VH and VL scores of the humanized antibodies, with a threshold set at 0.8. All VH and VL chains of our humanized antibodies exceed this threshold. Note that 2B04-hAb1.2 and 2B04-hAb1.3 are so similar in these metrics that they nearly overlap. The grey cross represents the mouse antibody, whose VH and VL scores are noticeably lower than the threshold. **c**, The histogram illustrates the human germline identity of the VH and VL regions for the parental mouse antibody (represented by the grey bar) and the five humanized antibodies (represented by the bars in other colors). Higher values indicate greater similarity to the closest human germline sequences. **d**, BLI kinetic analysis for the parental mouse antibody (grey) and five humanized antibodies: hAb1.1 (orange), hAb1.2 (green), hAb1.3 (purple), hAb1.4 (blue) and hAb1.5 (light cyan) . The values of k_on_ (association rate) and k_off_ (dissociation rate) are also presented. The binding affinity K_D_ values are 0.12 nM, 0.27 nM, 0.15 nM, 0.40 nM, 1.03 nM, and 6.37 nM, respectively. **e**, The 3D perspective illustrations are presented for the number and positions of mutations in each humanized antibody. **f**, The left panel displays the crystal structure of the parental mouse antibody, depicting a hydrogen bond between the residue LYS71 of FR3 and the residue THR53 of CDR2. The right panel shows predicted structures of the humanized antibodies, indicating that the residue ARG71 of FR3 and the residue THR53 of CDR2 still maintain the hydrogen bond. **g**, This 3D perspective illustrations shows that the residue VAL71 in FR3 and the residue THR53 in CDR2 of the predicted structures (hAb1.4 and hAb1.5) do not form a hydrogen bond.

All selected humanized antibodies and the murine antibody 2B04 are produced in Human Embryonic Kidney Cells 293 (HEK-293), with details of the purification and binding affinity experiments described in the Methods section. Initially, ELISA experiments confirm that all humanized antibodies retain their antigen-binding ability (see Supplementary Table S1). Furthermore, Bio-Layer interferometry (BLI) results (see Figure 3d) show that our humanized antibody hAb1.2 exhibits comparable binding affinity (K_D_) to 2B04 (0.15 nM for hAb1.2 versus 0.12 nM for mAb 2B04), indicating that the affinity is maintained despite 39 total mutation residues (see Figure 3e). Other two humanized antibodies, hAb1.1 and hAb1.3, retain similar binding affinities, with values of 0.27 nM and 0.40 nM, respectively, as shown in Figure 3d. Additionally, while the binding affinities of hAb1.4 and hAb1.5 decreased by one order of magnitude, they remained within the nanomolar range.

To better understand the structural differences that lead to variations in binding affinity among these humanized antibodies, we use AlphaFold3 to predict their 3D structures. The panels in Figure 3e displays the positions of mutation residues from a structural perspective for all humanized antibodies (sequence-level analysis is provided in Supplementary Figure S5). Based on previous research ^61^, which suggests that most interactions between the SARS-CoV-2 RBD and mAb 2B04 arise from the heavy chain, we focus on exploring the differences among the heavy chains of these humanized antibodies. In mAb 2B04, a hydrogen bond forms between LYS71 in FR3 and THR53 in CDR2 (see the left panel of Figure 3f). In hAb1.1, hAb1.2, and hAb1.3, the mutation of LYS71 to ARG preserves this hydrogen bond with THR53 (see the right panel of Figure 3f). However, in hAb1.4 and hAb1.5, LYS71 is mutated to VAL, which does not form a hydrogen bond with THR53 (see Figure 3g). This suggests that the hydrogen bond contributes to the stabilization of CDR1, CDR2, and CDR3, helping maintain a conformation that closely aligns with that of mAb 2B04.

### 2.3. Evaluation of humanness performance and experimental validation of HuDiff-Nb

#### Humanization process and performance of HuDiff-Nb

As shown in Figure 4a, HuDiff-Nb employs two distinct sampling processes. The first method, labeled as ‘de’, humanizes the nanobody from scratch based only on the input of the three complementarity-determining regions (CDRs) from the VHH domain (single heavy-chain variable domain) sequences, similar to the process used for HuDiff-Ab. The second method, labeled as ‘inp’, addresses a unique characteristic of nanobodies: the absence of a light chain. This absence leads to a higher proportion of hydrophilic residues in the FR2 region, which has been shown to significantly influence nanobody expression and stability ^62,63^. To accommodate this, we introduce the inpainting method that preserves these critical residues during the humanization process.

**Figure 4.**
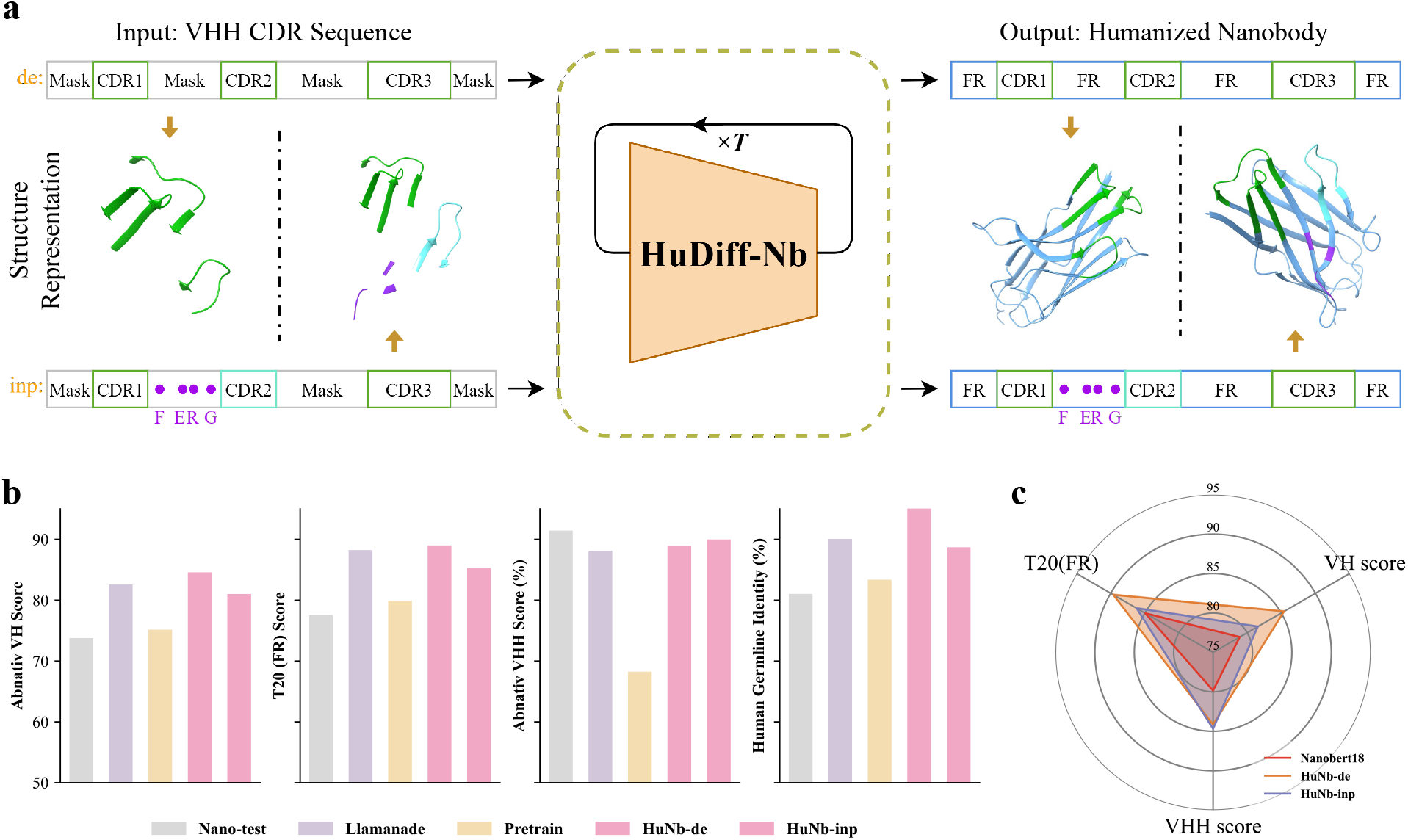
Humanization process and performance of HuDiff-Nb model. **a**, The diagram outlines two humanization processes of the HuDiff-Nb model. ‘de’ represents the process of humanizing nanobodies from scratch, where only the three CDR sequences (green blocks) of the VHH chain are used as input. After T sampling iterations, a humanized nanobody is obtained, as depicted by the structural representation showing the combination of the CDR structure (green cartoon) with the FR structure (blue cartoon). ‘inp’ represents a process where key residues (purple amino acid abbreviations) in the FR2 sequence are fixed, inspired by the Llamanade approach using their definitions of CDR2 (light blue block). The humanization process is performed similarly to the ‘de’ method, resulting in a humanized nanobody, as shown in the structural representation, where the fixed residues are combined with the FR residues (blue cartoon). **b**, The grey bars depict the performance of the Nano300 test dataset across various metrics, while the colored bars illustrate the performance of humanized nanobodies generated by the Llamanade pipeline and our methods (Pretrain, HuNb-de, HuNb-inp) on the same metrics within the Nano300 test dataset. The Abnativ VH score, T20 FR score, and Human germline identity are evaluate the humanness of humanized or original nanobodies. The Abnativ VHH score assesses the VHH nativeness of both the nanobodies themselves and the humanized nanobodies. **c**, Comparison of the humanness performance of our humanized nanobodies using different sampling methods with that of experimentally humanized nanobodies in the Nanobert18 humanized dataset.

To benchmark our method, we select the Llamanade approach ^39^ for comparison. The humanization performance of nanobodies in previous AbNativ experiments ^40^ is evaluated using two key metrics: the AbNativ humanness score and the AbNativ VHH-nativeness score, which reflect how closely a nanobody resembles human antibodies while preserving essential camelid features. Consequently, to ensure comprehensive assessment, we incorporated these two metrics, along with T20 score of the framework region and human germline identity. To evaluate both methods, we compiled two datasets: Nanobert18, containing only humanized nanobodies (excluding parental nanobodies), and Nano300, which includes only unhumanized nanobodies (see Methods for details).

We initially compare the performance of HuDiff-Nb against the Llamanade method using the Nano300 dataset. As shown in Figure 4b, while the raw nanobodies (Nano-test) exhibit low humanness and high VHH-nativeness, both the Llamanade and HuDiff-Nb humanized nanobodies show significant improvements in humanness, indicated by higher VH scores, T20 FR scores, and Human Germline identity. Notably, HuDiff-Nb, through both the ‘de’ and ‘inp’ methods, outperforms the Llamanade method in terms of humanness. Interestingly, despite ‘de’ sampling method of HuDiff-Nb (denoted as HuNb-de) not being fine-tuned using the T20 score, it achieves the higher T20 FR score compared to the Llamanade, which specifically uses T20 as a humanness filter. Additionally, we observe that the VH score of the pre-trained version of HuDiff-Nb (denoted as Pretrain) is lower than that of both HuNb-de and HuNb-inp (HuDiff-Nb ‘inp’ method), and its AbNativ VHH score decreases significantly compared to the raw AbNativ VHH score from the Nano300 test dataset (see Figure 4b). This result aligns with expectations, suggesting an out-of-distribution issue for CDR sequences between VHH and VH.

We further validate the humanization performance of HuDiff-Nb using the Nanobert18 experimental dataset. As shown in Figure 4c, both HuNb-de and HuNb-inp achieve higher T20 FR and AbNativ VH scores compared to the experimental humanized nanobodies, while also maintaining a higher AbNativ VHH score. However, since the test dataset lacks the parental nanobody sequences corresponding to the humanized nanobodies, we are unable to evaluate the performance of the Llamanade method, as its humanization process relies on the FR sequences of the parental nanobodies.

#### Humanization of nanobody 3-2A2-4 targeting SARS-CoV-2 RBD and experimental Validation

The crystal structure of wild-type nanobody 3-2A2-4, which binds specifically to SARS-CoV-2 receptor-binding domain (RBD) with an IC50 value of 0.046 μg/ml ^54^, has been experimentally determined. To assess the efficacy of HuDiff-Nb in humanizing nanobodies, in silico humanized variants of the wild-type nanobody are generated using two sampling methods (‘de’ and ‘inp’), from which five nanobodies are selected for production and subsequent in vitro characterization. Among the selected humanized variants, three (hNb1.1, hNb1.2, hNb1.3) are obtained using the ‘de’ sampling method, while two (hNb1.4, hNb1.5) are generated using the ‘inp’ sampling method. Notably, the FR T20 scores of these humanized nanobodies exceed those of 3-2A2-4 (see Figure 5a). As anticipated, AbNativ Humanness score (VH score) and AbNativ VHH-nativeness score of all humanized nanobodies are higher than those of 3-2A2-4 (see Figure 5b). We observe that the humanness scores of the humanized nanobodies derived from the ‘de’ method are higher than those derived from the ‘inp’ method. Furthermore, we calculate the human germline identity and FR2 distance between the humanized nanobodies and 3-2A2-4. All humanized nanobodies exhibit higher human germline identity than 3-2A2-4 (see Figure 5c), indicating that the FR sequences are closer to those of humans. As shown in Figure 5d, FR2 distance, which measures the number of mutated residues in the FR2 sequences, is greater for the humanized nanobodies (hNb1.1, hNb1.2, hNb1.3) using the ‘de’ method compared to those humanized nanobodies (hNb1.4, hNb1.5) with the ‘inp’ method.

**Figure 5.**
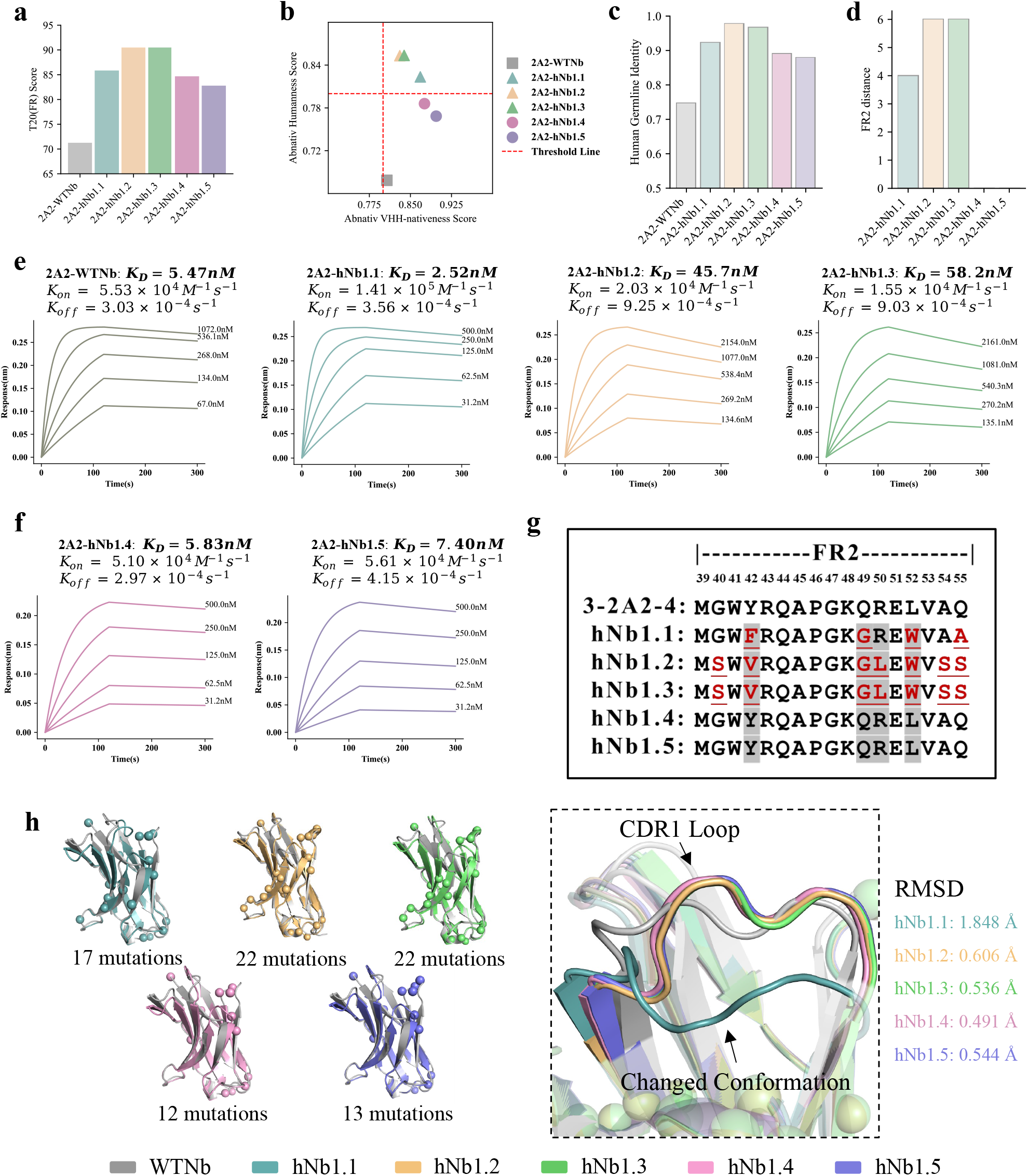
Humanness and binding affinity of humanized nanobodies. **a**, Comparison of T20 scores between the humanized nanobodies framework regions (FR) and wild-type 3-2A2-4. All T20 scores of the humanized nanobodies are higher than that of the wild-type 3-2A2-4. **b**, The X-axis represents VHH-nativeness, and the Y-axis indicates humanness (VH score). The threshold line is set at 0.8. The grey square represents the wild-type 3-2A2-4, whose humanness is below the threshold. Colored triangles represent nanobodies humanized using the ‘de’ method, while colored circles indicate nanobodies humanized using the ‘inp’ method. **c**, The bar plot illustrates the human germline identity of the wild-type 3-2A2-4 and the humanized nanobodies. The humanized nanobodies exhibit a higher human germline identity compared to the wild-type nanobody. **d**, The bar plot illustrates the FR2 distance between the humanized nanobodies and the wild-type nanobody. The FR2 distance of the humanized nanobodies generated by the ‘de’ sampling method is either 4 or 6, whereas the FR2 distance of the humanized nanobodies produced by the ‘inp’ method is 0. **e**, BLI kinetic analysis of the wild-type 3-2A2-4 and the humanized nanobodies (hNb1.1 in baby blue, hNb1.2 in yellow, and hNb1.3 in light green) generated by the ‘de’ method reveals binding affinities of 5.47 nM, 2.52 nM, 45.7 nM, and 58.2 nM, respectively. **f**, BLI kinetic analysis of the humanized nanobodies (hNb1.4 in pink, hNb1.5 in modena) generated by the ‘inp’ method reveals binding affinities of 5.83 nM and 7.40 nM, respectively. **g**, The FR2 alignment of the wild-type nanobody and the humanized nanobodies is shown, with the grey-highlighted text marking the positions of key residues that must be preserved using the ‘inp’ method. **h**, The left panel illustrates the number of mutations in the humanized nanobodies and their positions in 3D perspective. The right panel demonstrates the conformational changes in CDR1 of hNb1.1 compared to the wild-type nanobody and the other humanized nanobodies.

All humanized nanobodies and 3-2A2-4 are produced in Human Embryonic Kidney Cells 293 (HEK-293), with the purification process detailed in the Methods section. We first validate whether the humanized nanobodies retain their antigen-binding capability through ELISA experiments (see Supplementary Table S2). Following this, we assess the binding affinities by measuring the dissociation constants K_D_ values for all selected humanized nanobodies and 3-2A2-4 in their interactions with SARS-CoV-2 RBD using BLI. BLI experiment demonstrates that the K_D_ value for wild-type 3-2A2-4 is 5.47 nM (see Figure 5e). Notably, one of the five humanized nanobodies, hNb1.1, exhibits a K_D_ value of 2.52 nM, representing a 54% improvement in binding affinity compared to 3-2A2-4. Two others, hNb1.4 and hNb1.5, display comparable or slightly reduced binding affinities relative to the wild-type nanobody (see Figure 5f). These results suggest that HuDiff-Nb effectively humanizes the nanobody while maintaining or enhancing its binding affinity compared to the parental nanobody. Furthermore, it indicates that ‘de’ sampling method of HuDiff-Nb may have great potential for generating improved, better-humanized nanobodies.

To investigate why hNb1.1 exhibits greater binding affinity than the wild-type nanobody, we conducted a structural analysis of their 3D conformations. The 3D structures of all humanized nanobodies and wild-type nanobody are predicted using AlphaFold3 web server. As shown in Figure 5h, the left panel highlights the mutation positions of all humanized nanobodies from a structural perspective (the sequence-level analysis is available in Supplementary Figure S6), revealing that humanized nanobodies generated by the ‘de’ sampling method have more mutations than those from the ‘inp’ method. Upon close comparison of the structures, we observe no significant conformational differences in CDR2 and CDR3, except for CDR1. The right panel of Figure 5h shows the CDR1 conformations of all humanized nanobodies, where the conformation of hNb1.1 deviates significantly from the others, with a root mean square deviation (RMSD) of 1.848 Å compared to the 3-2A2-4. This conformational change likely contributes to its enhanced binding affinity. Additionally, through sequence alignment of the FR2, we aim to uncover the factors contributing to the differences in binding affinities among the humanized nanobodies. As illustrated in Figure 5g, hNb1.2 and hNb1.3 exhibit notable residue differences compared to hNb1.1, hNb1.4, and hNb1.5. These variations suggest that mutations at residues 40, 42, 50, 54, and 55 are crucial for maintaining the binding affinity of 3-2A2-4. Notably, we observe that the conserved residue at position 42 (shaded in gray) in hNb1.1 mutates to phenylalanine (Phe, characterized by a benzene ring as its R group), while hNb1.4 and hNb1.5, derived via the ‘inp’ method, retain the amino acid tyrosine (Tyr, containing a phenol group). Compared to Tyr, Phe side chain lacks an additional hydroxyl group, resulting in a stronger hydrophobic effect. This enhanced hydrophobicity may explain why hNb1.1 demonstrates stronger affinity than hNb1.4, hNb1.5, and even 3-2A2-4. This finding highlight the importance of specific residues, such as those in the FR2, in preserving the functional integrity of nanobodies during the humanization process.

## 3. Discussion

In this work, we present an efficient and adaptive diffusion approach called HuDiff, specifically designed for the effective humanization of antibodies and nanobodies. By applying this approach to various target sequences, we develop HuDiff-Ab for antibodies and HuDiff-Nb for nanobodies. Both of them share a similar architecture, consisting of encoders, hidden blocks, and a decoder. Besides, our approach requires only the CDR sequences during the humanization process, eliminating the need for a human template. In tests across different datasets, the humanization performance of HuDiff-Ab is comparable to that of the Sapiens, and even surpasses it in some metrics. For nanobody humanization, we develop a new sampling method that considers key residues impacting nanobody expression, which is distinct from antibody humanization. In various nanobody evaluation datasets, HuDiff-Nb not only improves the humanness of humanized nanobodies but also preserves their VHH-nativeness. To validate that binding affinity is maintained, the antibody 2B04 and nanobody 3-2A2-4 are selected, both targeting the SARS-CoV-2 RBD antigen. HuDiff-Ab and HuDiff-Nb are successfully used to humanize these sequences and generate five highly humanized variants of each. In subsequent BLI experiments, all humanized sequences preserve their ability to bind to the target. Notably, the top humanized antibody exhibited binding affinity nearly equivalent to that of the parental mouse antibody 2B04, while the top humanized nanobody unexpectedly demonstrated improved binding affinity compared to the parental alpaca nanobody 3-2A2-4. These findings confirm that our training approach is effective for humanization tasks, achieving high humanness while preserving, and enhancing binding affinity in some cases.

In the task of mouse antibody humanization, our approach uniquely leverages paired antibody sequences to train HuDiff-Ab, distinguishing it from other humanization methods. These paired sequences are aligned using the IMGT numbering scheme. Although our training dataset includes approximately 2 million paired sequences, significantly fewer than Sapiens’ 20 million heavy chains and 19 million light chains, the humanness performance of our model remains comparable to Sapiens. Furthermore, when combined with other evaluation methods, HuDiff-Ab demonstrates even higher humanness performance, as indicated by metrics such as the OASis and T20 scores. These findings suggest that paired sequences may offer significant advantages for antibody tasks, consistent with observations from other studies ^56^. Additionally, we explore the effectiveness of HuDiff-Ab on the Putative dataset, as shown in Supplementary Table S3, where the humanness performance of HuDiff-Ab closely aligns with the true experimental results and surpasses that of the Sapiens1. Additionally, we explore the impact of sampling order, given that our method is based on the autoregressive diffusion model ^64^, which allows the pretrained model to sample sequences in a left-to-right manner. As shown in Supplementary Figure S7, the performance remains consistent between shuffled order and left-to-right sampling.

In the camel nanobody humanization task, our two-step training strategy proves beneficial to HuDiff-Nb as it addresses the out-of-distribution challenge between VH and VHH CDR sequences. HuDiff-Nb first undergoes pre-training on heavy chains of conventional antibodies, and then fine-tuning to incorporate CDR information from VHH sequences. To our knowledge, no other models or methods directly account for nanobody CDR sequences during the humanization training phase. Building on this strategy, we explore an alternative humanization approach that focuses solely on optimizing the VH score, without considering the unique native properties of nanobodies. In contrast, our primary objective integrates both VH score and VHH nativeness score during fine-tuning. Our ablation study (see Supplementary Figure S8a, b) demonstrates that the version optimizing only the VH score (referred to as HuNb2) still achieves excellent performance in both humanization and VHH nativeness, similar to our primary objective (referred to as HuNb1).

Drawing on insights from recent studies ^62,63,65^, it is evident that the absence of a VL domain in nanobodies leads to an increased presence of hydrophilic amino acids (e.g., Phe42, Glu49, Arg50, and Gly52) in the FR2 region, in contrast to conventional VH regions, which are enriched with hydrophobic residues such as Val42, Gly49, Leu50, and Trp52. This change enhances the hydrophilic surface of VHHs and contributes significantly to nanobody stability. Consequently, our ‘inp’ sampling method for humanizing nanobodies preserves essential residues at key positions, supporting both stability and expression. Moreover, unlike the approaches of Llamanade and AbNativ, which incorporate a structure prediction stage, our method works exclusively at the sequence level. This allows for a more efficient process, maintaining the quality of humanization while being faster and more streamlined.

We find that employing HuDiff-Nb for large-scale humanization of nanobody 3-2A2-4 generates approximately one thousand distinct humanized nanobodies per sampling approach. The limited number of humanized nanobodies produced per approach likely stems from two key factors. Firstly, there is a high degree of similarity between the FR sequences of nanobody and antibody heavy chain. Research ^62,39^ highlights that although certain residues differ between FR2 and FR3 of VH and VHH, these differences are not extensive. The second factor involves the use of our auxiliary models, AbNativ VH and AbNativ VHH. During fine-tuning, these models refine the training objectives, potentially helping HuDiff-Nb more accurately capture the specific interactions between the FR sequences of nanobody and antibody heavy chains, thus reducing uncertainty in the diffusion process. Additionally, although previous studies ^7,66^ suggested that a uniform residue framework might suffice for replacing the FR of an unhumanized nanobody, the differing binding affinities observed in HuDiff-Nb’s experimental results underscore the need for exploring various humanization strategies. Both the limited evidence from past studies ^7,66^ and our own findings indicate that a customized approach to nanobody humanization is critical. This tailored method not only preserves the original binding affinity but also has the potential to enhance it.

Our approach humanizes antibodies and nanobodies from scratch, generating FR sequences based on CDR sequences, which differs from conventional humanization methods that require back mutations after grafting. This template-free method enables the generation of a diverse range of humanized FR sequences. As shown in Table 1, HuDiff-Ab can be combined with other evaluation methods to select the most effective humanized sequences from this diversity. Additionally, if the parent FR sequence of a nanobody is unknown, HuDiff-Nb can humanize it by inputting only the CDR residues, offering a more convenient alternative than methods like Llamanade, which require the entire sequence.

Overall, our approach demonstrates strong humanization capabilities for both antibodies and nanobodies, as evidenced by significant improvements in humanness compared to parental mouse antibodies and alpaca nanobodies. Importantly, the binding affinities of humanized sequences are maintained at comparable levels to their parental counterparts, and in some cases, even greatly improved. Through structural analysis, we find the humanized sequences retained conformations of the complementarity-determining regions (CDRs) that are close to those of the parental sequences, preserving CDR structural integrity and stabilizing the interactions between them. This structural stability likely contributes to the retention of binding affinity. In contrast, while structural similarity is important, it may also be beneficial to assess how conformational changes affect function. For instance, some structural changes can enhance binding affinity, as shown in Figure 5f, while others may reduce it, as demonstrated in Supplementary Figure S9. Considering all these factors could further enhance the humanization process.

In summary, our method successfully implements the humanization process for both antibodies and nanobodies, with its performance validated through both in silico and wet lab experiments. We anticipate that our approach, HuDiff, will serve as a valuable tool for researchers advancing antibody and nanobody humanization. Looking ahead, based on our structural analysis, we believe that integrating structural information during the training phase of future humanization models may benefit antibody humanization design, though this remains uncertain. While incorporating structural information in the humanization process, as seen in methods like Abnativ and Llamanade, is effective, it may not be efficient. Our approach marks a significant step forward in the development of next-generation humanization techniques.

## 4. Methods

### Training dataset and preprocessing

#### Antibody data preprocessing

For the HuDiff-Ab, the pre-training phase utilizes a paired human antibody dataset sourced from Observed Antibody Space (OAS) database ^67,57^. In the fine-tuning phase, we collect a paired mouse antibody dataset from OAS, PLAbDAb ^68^, and ABSD^1^ database. The pre-training dataset contains nearly two million paired sequences (1,954,079), while the fine-tuning dataset consists of approximately twenty thousand paired sequences. We filter out sequences missing Framework Region 1 (FR1) or containing unknown residues, and remove duplicates. After this curation, the pre-training dataset comprises 1,738,321 paired sequences, which is split into training and validation sets at a 19:1 ratio. The fine-tuning dataset, consisting of 19,387 paired sequences, is divided following the same ratio.

#### Nanobody data preprocessing

The training process of HuDiff-Nb is also divided into two distinct phases: pre-training and fine-tuning, each utilizing its own specific dataset. For the pre-training phase, we compile an unpaired heavy chain dataset from human sequences in the OAS database. From this database, we randomly select over three million heavy chain sequences, ensuring the inclusion of the standard HV1 to HV7 germline families. In the fine-tuning phase, we download all available camelid sequences from the OAS database. After applying the same filtering criteria used for HuDiff-Ab, we obtain a pre-training dataset of 3,395,594 sequences and a fine-tuning dataset of 841,488 sequences to train the HuDiff-Nb. Both the unpaired heavy chain and single-domain heavy chain training datasets are aligned using the IMGT numbering scheme ^59^. Additionally, during the fine-tuning stage, we re-encode each VHH sequence from the camelid dataset using the AHO numbering scheme. This step is necessary to facilitate the conversion between numbering systems and to further refine the fine-tuning process of HuDiff-Nb.

### Benchmark dataset

Humab25 dataset comprises 25 experimentally validated paired antibodies derived from the Humab ^35^. Additionally, we compile HuAb348 dataset from patent records, which comprises 348 experimentally validated humanized paired antibodies and their corresponding parental mouse antibodies. These datasets serve as test sets to evaluate our model’s performance against Sapiens and traditional methods, as discussed in the main text. The reconstructed Putative152 dataset, where mouse sequences are identified by sequence similarity, is treated as an additional test set due to concerns about the authenticity of the data, with its humanness performance provided in Supplementary Table S3. For nanobodies, we compile a test dataset, Nanobert18, consisting of 18 humanized nanobodies sourced from the publication by Hadsund et al. (2024), without including the parental non-humanized nanobodies. To further evaluate and benchmark nanobody humanization performance against Llamanade, we construct a new test set, Nano300, consisting of 300 nanobodies that are not included in our training dataset.

### Baseline

**Sapiens**: Sapiens model is obtained from BioPhi GitHub repository^2^. For HuAb348 dataset, we follow the humanization steps outlined in their paper. The performance results for the Humab25 dataset come directly from the BioPhi published paper ^37^. **Llamanade**: We install the Llamanade method from the Llamanade GitHub repository^3^, which is a computational pipeline designed to perform the humanization process for nanobodies. We follow the process described in their paper, using nanobody FASTA files for humanization. This process includes a structure prediction step, for which we use the one of default tools, Modeller ^69^, to predict the structure.

### Germline identity

In the study of germline identity, the “Graft CDRs onto Human Germline” function within the Abnumber tool ^70^ is employed to transplant CDRs from a given chain onto the closest matching human germline sequence. Consequently, germline identity is a metric we employ to quantify the degree of resemblance between the framework region sequence we have engineered and the most similar human germline sequence. The formula of the germline identity is:

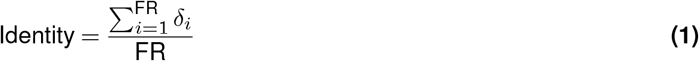

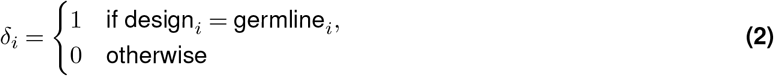

FR is the length of the framework region, which differs between the heavy and light chains. The design_*i*_ and germline_*i*_ represent the residues at corresponding positions in the alignment of the two sequences. This metric can be used to measure the rationality of sequences that we design from scratch.

### FR2 distance

We employ the Levenshtein distance ^71^, a string metric that quantifies the difference between two sequences, to calculate the FR2 distance. Specifically, Levenshtein distance represents the minimum number of single-character edits—insertions, deletions, or substitutions—required to transform one sequence into another. This metric provides a robust measure for assessing the degree of variation between FR2 sequences in our analysis.

### Gumbel-Softmax

Gumbel-Softmax ^72^ technique is employed to generate one-hot vectors from the output of pretrained model, as the Argmax function results in non-differentiable gradients. In detail, we introduce Gumbel noise to each output logit of the pretrained model. This noise is generated by applying the logarithm twice to a uniform distribution, a method similar to the reparameterization trick where normally distributed noise is added to the mean. After adding Gumbel noise, we apply a temperature parameter (set to 1 in our case) to control the noise logits. We then apply the Softmax function to calculate the probabilities of these noise logits. Then, non-differentiable values are added to obtain the one-hot vector. With this approach, once we obtain the humanness score of a residue using AbNativ ^40^, the gradient can be backpropagated, allowing fine-tuning of HuDiff-Ab and HuDiff-Nb.

### Architectures of HuDiff models

#### HuDiff-Ab

HuDiff-Ab is designed to accept paired and aligned heavy-chain and light-chain sequences as input. We employ the IMGT numbering scheme to align these sequences, resulting in a heavy chain length of 152 and a light chain length of 139. Consequently, the total length of the input sequence is 291. We use the symbol ‘-’ to denote padding tokens, ‘#’ to denote diffused state tokens, and ‘X’ for unknown residue types. Along with the standard 20 residues, the token size is brought to 23. Once the tokenizer processes the string of input aligned residues, these string residues are converted into a numerical vector. HuDiff-Ab is composed of several components: a sequence encoder, an extra information encoder, a dual CNN Block, a Roformer, and a decoder. The sequence encoder initially employs an Embedding layer to embed the diffused numerical vector of the paired sequence X_T_. This embedded tensor is then encoded using the Dual CNN Block of the sequence encoder. The Dual CNN Block consists of two parallel layers - heavy and light - each designed to analyze the features of the heavy and light sequences, respectively. However, solely learning the sequence features is insufficient to fully represent the sequence information. To address this, we use an extra information encoder, which includes a position block, a region block, and a chain block. These blocks encode the residue position, CDR and FR region index, and chain type information, respectively. For the position block, we use sine-cosine position encoding for each residue position and an MLP network to map the position information. The region block employs a region embedding to embed the region vector. Given that a chain can be divided into three CDR regions and four FR regions, our region embedding’s maximum index in the region vector is 6. The chain block aids the model in determining which part of the sequence belongs to the heavy chain, and which part belongs to the light chain (*λ* or *κ*). We ensure that the output tensors of sequence encoder and extra information encoder have the same dimension, and then we add and concatenate them. This provides a comprehensive representation of the paired heavy-chain and light-chain sequences to the HuDiff-Ab. We then use an independent Dual CNN block, similar in architecture to the previous one but with fewer layers, to explore the paired antibody information beyond just the sequence itself. Following this, we employ a self-attention block, constructed from several self-attention layers and introducing the Rotary Position Embedding (ROPE), to capture sequence dependencies. This is done so that the model can capture the relationship among various regions of the sequences. Finally, we employ a decoder, implemented as a MLP, to generate the logits for the predicted sequence.

PyTorch version 1.13 is used to construct HuDiff-Ab. The kernel size for all CNN blocks is set to 7. Training is conducted using the Adam optimizer with an initial learning rate of 1.e-4, with a batch size of 256. We employ a ReduceLROnPlateau scheduler with patience of 10 and a decay factor of 0.6 to adjust the learning rate.

#### Objective function

In the traditional autoregressive model, the distribution is factorized through the chain rule 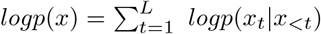, following a pre-determined decoding order from left to right. In contrast, the training process of HuDiff-Ab, built using the autoregressive diffusion method^64^, mirrors the absorbing state concept in D3PM, as described in the research ^73^. While both aim to recover a target token from a masked state, D3PM constructs a transition matrix, whereas HuDiff-Ab corrupts the sequence directly, resembling Masked Language Model ^74^. During pre-training, a random sequence *σ* of framework region residues is uniformly drawn from *S*_*F*_, the set of all possible permutations of the integers from 1 to *F* . The symbol *F* represents the FR length of human sequences. Using Jensen’s inequality, the log-likelihood for the order-agnostic autoregressive model can be expressed as:

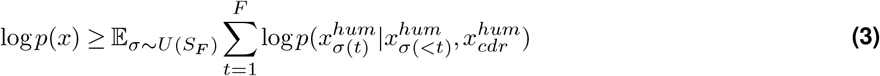

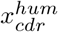 denotes the information contained within the CDR. As outlined by the research ^64^, an alternative objective to Equation 3 can be derived by replacing the summation over *t* with an expectation that is appropriately re-weighted:

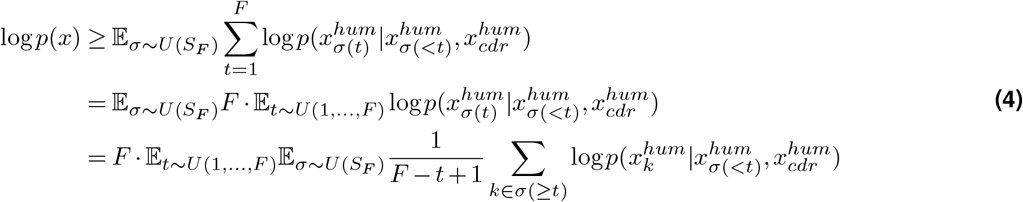

each loss at time step *t* can be regarded as *ℒ*_*t*_:

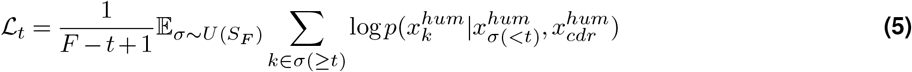

Consequently, we can write overall expected log likelihood log *p*(*x*) as:

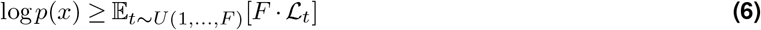

During our pre-training procedure, given that our input and output share a one-to-one correspondence, we incorporate an additional assistant loss. This loss ensures that CDR residues of the output sequences remain identical to CDR residues of the input sequences. Consequently, our training loss can be simply expressed as follows:

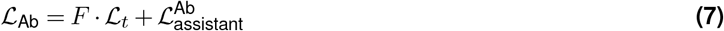

Here, the first *ℒ*_*t*_ can be interpreted as a reconstruction loss, which indicates that, when the diffused state *x*_*t*_ of the sequence is input, the model needs to predict the original state *x*_0_. During the fine-tuning procedure, the input to HuDiff-Ab consists of the diffused paired mouse antibodies, denoted as 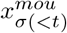, while the output comprises the logits of 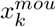. In this context, all residues *σ*(≥ *t*) are humanized based on our pre-training objective. To enhance the guidance of HuDiff-Ab training, 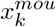 is transformed into a one-hot vector using Gumbel Softmax trick. The VH score for heavy chain residues and the VL score (*λ* or *κ*) for light chain residues are then evaluated through the AbNativ ^40^. The fine-tuning process can be formulated as follows:

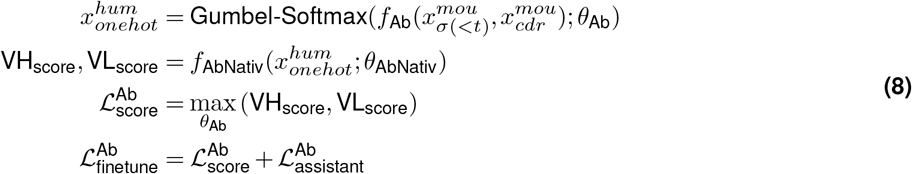

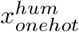 can be considered as a hypothetical representation of humanized antibodies. The VH_score_ and VL_score_ represent the scores generated by the AbNativ. We then fine-tune the HuDiff-Ab using the loss objective 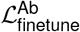 to enhance the humanness of the resulting humanized sequences.

#### Hudiff-Nb

The design of HuDiff-Nb bears resemblance to our previously developed HuDiff-Ab. Unlike HuDiff-Ab’s sequence encoder, HuDiff-Nb’s sequence encoder comprises a singular CNN Block tailored to its needs. This modification is because nanobody humanization focuses solely on a single chain, thus eliminating the requirement of chain block. Consequently, we only employ the position and region encoder to integrate the additional information into the HuDiff-Nb. All unpaired heavy chain data is preprocessed in a manner akin to the paired sequence data we handled previously. The notable distinction lies in the sequence length, which is now 152, reflecting the absence of the light chain. Following the integration and concatenation of the output features from both the sequence and region encoders, a CNN Block and a self-attention block are introduced to capture the hidden features of the chain. As with HuDiff-Ab, the training pipeline for HuDiff-Nb follows two phases. Firstly, we focus on a pre-training diffusion model exclusively designed for the heavy chain of a conventional antibody. The loss function employed in this learning phase is identical to the one used in the HuDiff-Ab, which is expressed as follows:

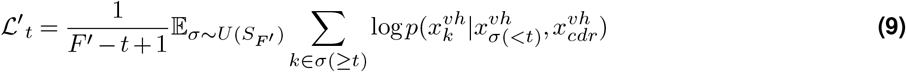

where *vh* represents the heavy chain of a conventional antibody. And *F* ^*′*^ represents the FR length of VH sequences. In the second phase, known as the fine-tuning stage, HuDiff-Nb receives various diffused time-step sequences derived from the VHH dataset as input. Since HuDiff-Nb was pretrained on the heavy chain of conventional antibodies, it tends to output the 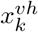 sequence, even when the input consists of a diffused VHH sequence. This behavior can be effectively represented by the conditional probability 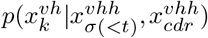. However, relying solely on the pretrained HuDiff-Nb presents a limitation: it is unable to accurately assess the generated residues within the context of the entire heavy chain sequence, particularly when accounting for the differences in CDR sequences between nanobodies and antibodies that fall outside the training distribution. To overcome this challenge, we incorporate the AbNativ ^40^ to evaluate the VH score (humanness score) and the VHH score (VHH-nativeness score) of the generated residues. We aim to utilize these scores to effectively guide the training process of HuDiff-Nb. Furthermore, our observations from wet lab experiments on humanized nanobodies developed using AbNativ indicate that these highly active humanized nanobodies not only achieve elevated humanness scores but also retain their VHH-native characteristics. Inspired by this finding, and in contrast to HuDiff-Ab which seeks to maximize both scores, we aim to optimize the training objective by maximizing the VH score while maintaining the changes in the VHH score (the difference in residue values between the humanized VHH sequences and their parental sequences). Additionally, akin to HuDiff-Ab, we utilize Gumbel Softmax technique to facilitate gradient backpropagation and preserve the integrity of the input and output CDR sequences, converting the output logits into one-hot vectors. We also incorporate the same auxiliary loss utilized during the pre-training stage. Ultimately, the fine-tuning loss function is simply expressed as follows:

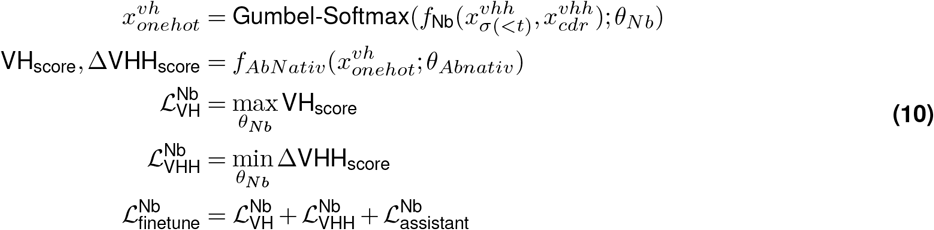

The sequence 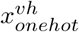 represents a hypothetical sequence of residues that, across different diffusion states of 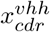 and 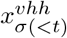, strives to mirror the framework sequence of a typical antibody heavy chain as closely as possible. This reflects the fact that the HuDiff-Nb’s initial training involved data from human heavy chains. To put it concisely, we have employed the humanization evaluations from AbNativ ^40^ to refine the HuDiff-Nb’s ability to capture the correlation between the heavy chain framework regions and CDR sequences of the VHH.

#### Structural modeling

In the research of structural modeling, AlphaFold3^48^ web server is used to directly predict the 3D structure of humanized antibodies and nanobodies. The server automatically selects the seed, and for each prediction, regardless of whether it pertains to antibodies or nanobodies, we only consider the top predicted conformation. Most regions of each top predicted structure exhibit high confidence (plDDT > 90), with the exception of the CDR regions, which generally show moderate confidence (70 < plDDT < 90). Notably, the CDR3 region often displays lower confidence (50 < plDDT < 70). The ipTM scores of the best predicted structures for the humanized antibodies are as follows: hAb1.1, hAb1.4, and hAb1.5 each have an ipTM score of 0.91; hAb1.2 has an ipTM score of 0.92; and hAb1.3 has an ipTM score of 0.89. Since nanobodies are single-chain, only pTM scores are available. We observe that hNb1.1, hNb1.2, and hNb1.3 each have a pTM score of 0.89, while hNb1.4 and hNb1.5 have pTM scores of 0.9.

### Production and purification of protein

#### Protein Production

The expression of humanized antibodies generated by HuDiff-Ab and humanized nanobodies generated by HuDiff-Nb begins with the amplification of the target gene through PCR. Notably, six His tags are appended to each humanized nanobody prior to Polymerase Chain Reaction (PCR) ^75^ to facilitate purification. Following digestion with restriction endonucleases, the gene is ligated into an expression vector and verified by sequencing. The confirmed plasmid is then transfected into competent HEK 293 cells for amplification. For antibodies and nanobodies expression, HEK 293 cells are cultured in serum-free CD medium and passaged three times before use. On the day of transfection, the cells are adjusted to a density of 2 × 10^6^ cells/mL in serum-free CD medium. The plasmid containing the target protein is mixed with the transfection reagent TF1 at a 1:10 (w/v) ratio and added to the cells, marking Day 0 of transfection. On Days 1, 3, and 5, serum-free 293 feed medium is added according to the reagent instructions. After the incubation period, the protein is purified from the cultured cells.

#### Protein purification

The purification of antibodies and nanobodies is accomplished using Protein A affinity chromatography. The procedure starts with the application of 50 mM Tris (pH 8.0) as the loading buffer, followed by two washing steps with 100 mM glycine (pH 3.0 and pH 2.7) and a stepwise elution gradient. SDS-PAGE analysis, performed under both reducing and non-reducing conditions, confirmed that the target antibodies or nanobodies are overexpressed in the HEK293 system.

### ELISA experiments

#### Antibody

SARS-CoV-2 RBD protein is coated onto the wells of a 96-well plate and incubated overnight at 4°C. The plate is then blocked with 2% BSA (bovine serum albumin) for 1 hour at room temperature to prevent non-specific binding. After washing with PBS-T (phosphate-buffered saline with 0.02% Tween 20), five humanized full-length antibodies are diluted to appropriate concentrations and added to the corresponding wells. The plate is incubated at room temperature for 1 hour. Following another wash, secondary antibodies, Rabbit Anti-Mouse IgG F(ab)2/HRP and Goat Anti-Human IgG (H+L)/HRP, are added and the plate is incubated for 1 hour at room temperature. After final washes, a color-developing solution is added, and the reaction is stopped by adding a stop solution (2 M sulfuric acid). The optical density (OD) is measured at 450 nm.

#### Nanobody

Five humanized VHH antibodies are coated onto a 96-well plate at a concentration of 1 μg/mL in 100 μL per well and incubated overnight at 4°C. The following day, the wells are blocked with 2% BSA (300 μL per well) for 1 hour at room temperature to minimize non-specific binding. After the blocking step, the plate is washed twice with 300 μL of wash buffer per well. The SARS-CoV-2 Spike RBD protein is then diluted to 0.5 μg/mL, and a negative control using sample diluent is prepared. Both the protein and the negative control are added to their respective wells (100 μL per well), mixed thoroughly, and incubated for 2 hours at room temperature. After incubation, the plate is washed three times with 300 μL of wash buffer per well, and Streptavidin/HRP secondary antibody, diluted to the appropriate concentration, is added to each well (100 μL per well) and incubated for 1 hour at room temperature. Following a final wash step (three washes), a color-developing solution is added to each well (200 μL per well) and incubated for 20 minutes at room temperature in the dark. The reaction is terminated by adding 50 μL of stop solution (2 M sulfuric acid) per well, and the optical density (OD) is measured at 450 nm.

#### BLI affinity experiments

Bio-Layer Interferometry (BLI) experiment is conducted using an Octet RED384 instrument and SA sensors (18-5019) from Sartorius. The running buffer is prepared as PBST (phosphate-buffered saline with Tween 20) supplemented with 0.1% BSA (bovine serum albumin). For the fixed ligand preparation, the SARS-CoV-2 (2019-nCoV) Spike RBD protein is diluted to 5 μg/ml using the running buffer and immobilized onto the SA sensors, achieving a binding level of approximately 0.6 nm. Different humanized antibodies are diluted to specific initial concentrations in the running buffer and subjected to a 2-fold serial dilution; this dilution process is also applied to various humanized nanobodies. The sensor with the immobilized Spike RBD is then placed in the BLI instrument, where the diluted humanized antibodies are sequentially introduced. The binding phase lasts for 120 seconds to facilitate interaction with the immobilized protein, followed by a dissociation phase of 180 seconds during which running buffer washes off unbound antibodies. The instrument records the wavelength shift data, which is subsequently analyzed using Data Analysis 12.0 software to determine the association rate constant (K_on_), dissociation rate constant (K_off_), and affinity constant (K_D_), thereby providing insights into the binding kinetics and affinity between the SARS-CoV-2 Spike RBD and humanized antibodies or nanobodies.

## Code and data Availability

All unprocessed sequences from the OAS database are available at https://opig.stats.ox.ac.uk/webapps/oas/. The PLABDAb database can be accessed at https://opig.stats.ox.ac.uk/webapps/plabdab/, and the ABSD database is available at https://absd.pasteur.cloud/about. All preprocessed datasets for training models, along with model checkpoints, are accessible at https://huggingface.co/cloud77/HuDiff. The code for our approach is available at https://github.com/TencentAI4S/HuDiff.

## Acknowledgements

We are grateful to “Tencent AI Lab Rhino-Bird Focused Research Program (Tencent AI Lab RBFR2023006, Tencent AI Lab RBFR2022006)” who provides the grants for this manuscript. We also thank the Supercomputing Center of Lanzhou University and Gansu Computation Center for supporting this paper.

## Author contributions

**Jian Ma:** Writing – review & editing, Writing – original draft, Visualization, Validation, Resources, Methodology, Investigation, Formal analysis, Data curation, Conceptualization. **Fandi Wu:** Writing – review & editing, Writing – original draft, Visualization, Validation, Supervision, Resources, Methodology, Investigation, Formal analysis, Data curation, Conceptualization. **Tingyang Xu:** Writing – review & editing, Visualization, Validation, Resources, Methodology, Investigation, Formal analysis, Data curation, Conceptualization. **Shaoyong Xu:** Writing – review & editing, Visualization, Validation, Resources, Methodology, Formal analysis, Data curation, Conceptualization. **Wei Liu:** Writing – review & editing, Visualization, Validation, Resources, Methodology, Formal analysis, Data curation. **Divin Yan:** Writing – review & editing, Visualization, Validation, Methodology, Formal analysis, Data curation. **Qifeng Bai:** Writing – review & editing, Visualization, Validation, Supervision, Resources, Project administration, Methodology, Investigation, Funding acquisition, Formal analysis, Data curation, Conceptualization. **Jianhua Yao:** Writing – review & editing, Visualization, Validation, Supervision, Resources, Project administration, Methodology, Investigation, Formal analysis, Data curation, Conceptualization.

## Supplementary

### Humanness scores

**OASis Score:** Derived from the OASis database, which uses a 9-per-peptide statistical frequency to construct its scores, OASis score delineates four tiers of antibody sequence humanization, each corresponding to specific frequency thresholds. Our evaluation mirrors the use of the OASis median value as a reference point, as it is utilized in the foundational study. **T20 Score:** This metric is calculated using a T20 score analyzer, which is inspired by the BLAST methodology, ranks sequences by their similarity to the query sequences. It then averages the identity percentages of top 20 matches to produce a T20 score. For the framework region, the process is similar, but it excludes the CDR sequences, focusing solely on the similarity within the framework region. **H Score:** This metric is calculated from the raw humanization score and its average. A higher H-score indicates a more human-like antibody, and the reverse is true for lower scores. **ABLSTM Score:** This metric is the output of a pre-trained LSTM model is trained on antibody sequences. Here, lower scores suggest a greater resemblance to naturally occurring antibodies. **AbNativ-humanness score and AbNativ-VHH-nativeness score:** The former is derived from models trained on human VH, VLambda, and VKappa chains, providing a metric for evaluating the similarity of a given chain to the learned distribution of human chains within the model. The latter, on the other hand, evaluates how similar a camelid chain is to the model’s learned distribution of camelid chains.

### In-depth Analysis of CDR Sequences in VH and VHH

We analyze the differences between the CDR sequences of VH and VHH in detail. Supplementary Figure S2 presents histograms that show the frequency of amino acid residues within the CDR sequences for both VH and VHH. Additionally, we provide the length distribution across various regions of the VH and VHH chains in Supplementary Figures S3 and S4. A detailed examination of the frequency data reveals a striking similarity in the amino acid composition of CDR1 for both VH and VHH (Supplementary Figure S2a), with a key difference at the terminal position: Tyrosine (Y) predominates in VH, whereas Cysteine (C) is more frequent in VHH. In CDR2, the second half exhibits comparable residue frequencies between VH and VHH (Supplementary Figure S2b), while slight differences emerge in the first half, especially at positions 57, 58, and 59, where ‘SPS’ is common in VH and ‘DSD’ is more frequent in VHH. CDR3, however, shows more pronounced differences in residue frequencies between VH and VHH, except for conserved positions at the beginning and end (Supplementary Figure S2c), underscoring a distinct divergence in residue composition within this region. Furthermore, when comparing the length variations in different regions of VH and VHH (see Supplementary Figures S3 and S4), significant differences arise in the CDR2 and CDR3 segments. The most common CDR2 lengths for VH and VHH are 7 and 8, respectively, though they appear in different proportions: 1:2 for VH and 1:1 for VHH. For CDR3, VH segments typically range from 10 to 20 residues, while VHH segments extend from 12 to 25 residues, indicating that CDR3 segments in VHHs tend to be longer than those in VHs.

**Figure S1.**
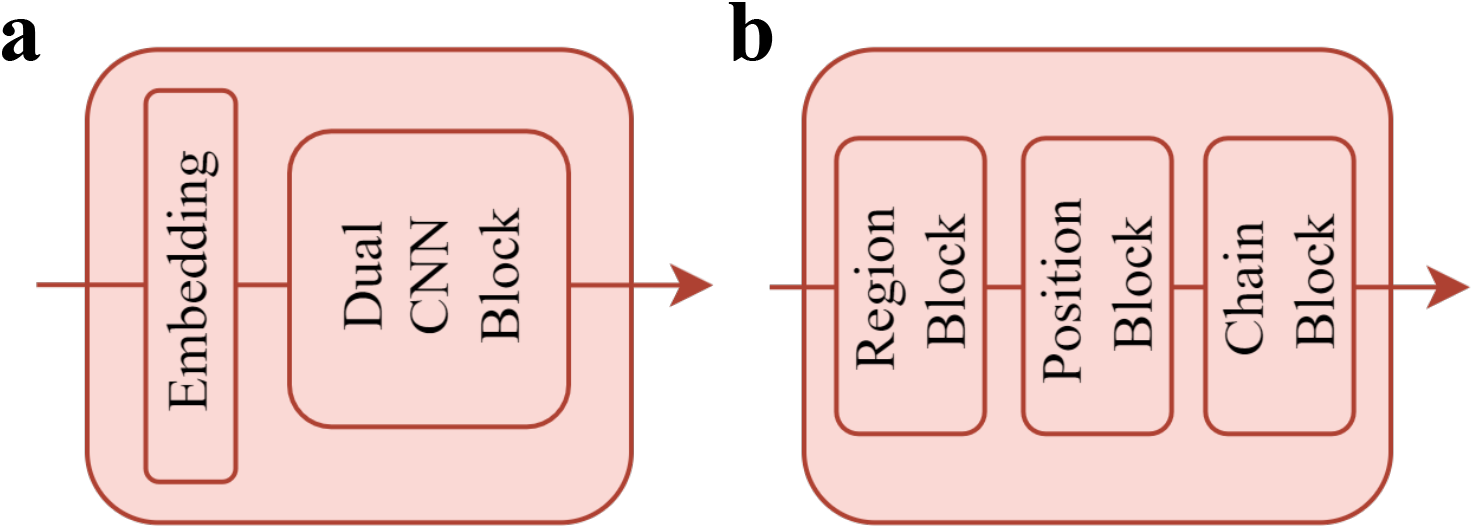
Architecture of the sequence encoder and information encoder. **a**, The sequence encoder includes Embedding and Dual CNN Block. **b**, The information encoder is composed of Region Block, Position Block, and Chain Block.

**Figure S2.**
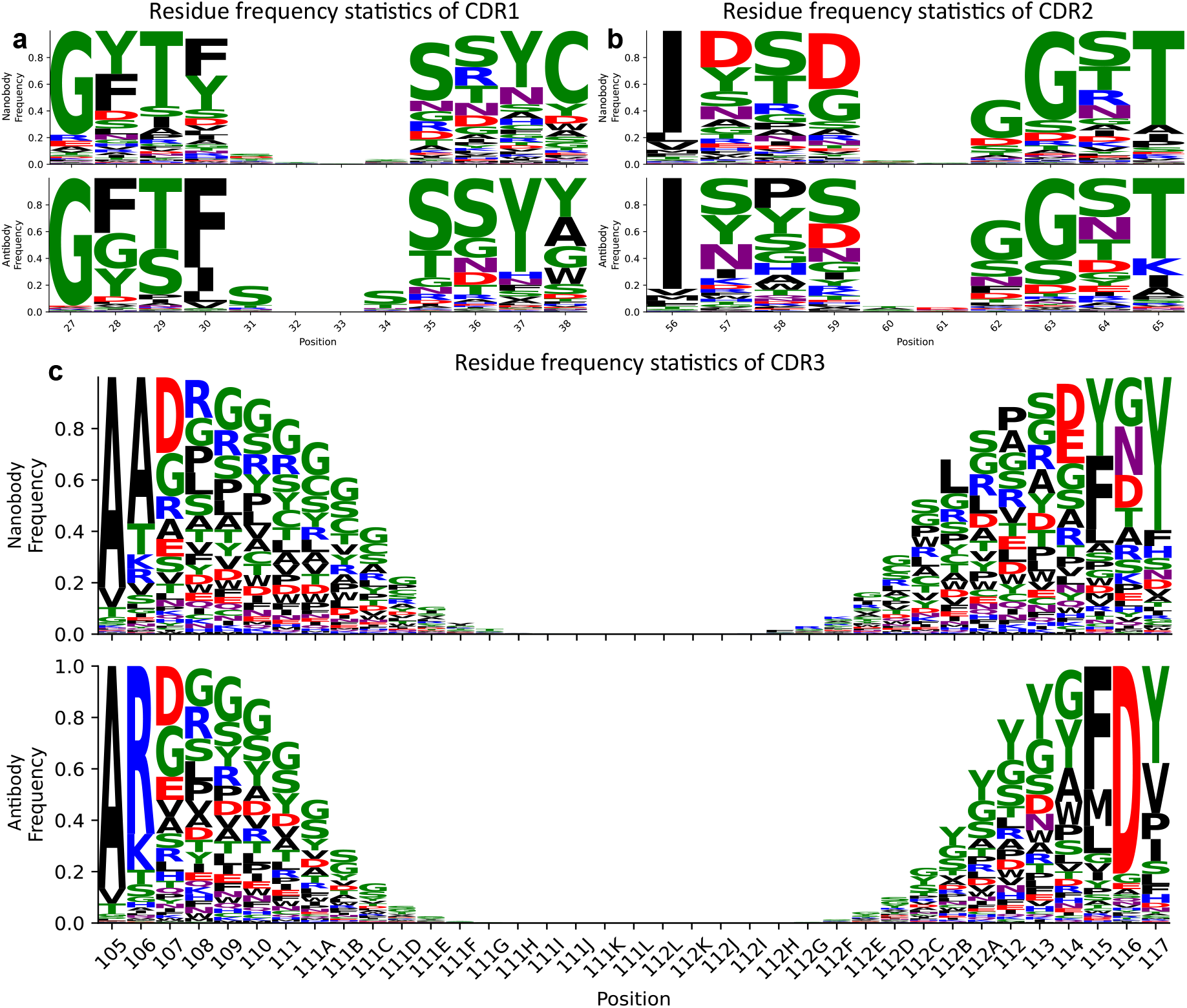
Comparison of CDR residues frequencies between heavy chain of antibodies and nanobodies. **a**, Comparison of aligned (IMGT) CDR1 residue frequencies between nanobodies and the heavy chain of antibodies. **b**, Comparison of aligned (IMGT) CDR2 residue frequencies between nanobodies and the heavy chain of antibodies. **c**, Comparison of aligned (IMGT) CDR3 residue frequencies between nanobodies and the heavy chain of antibodies.

**Figure S3.**
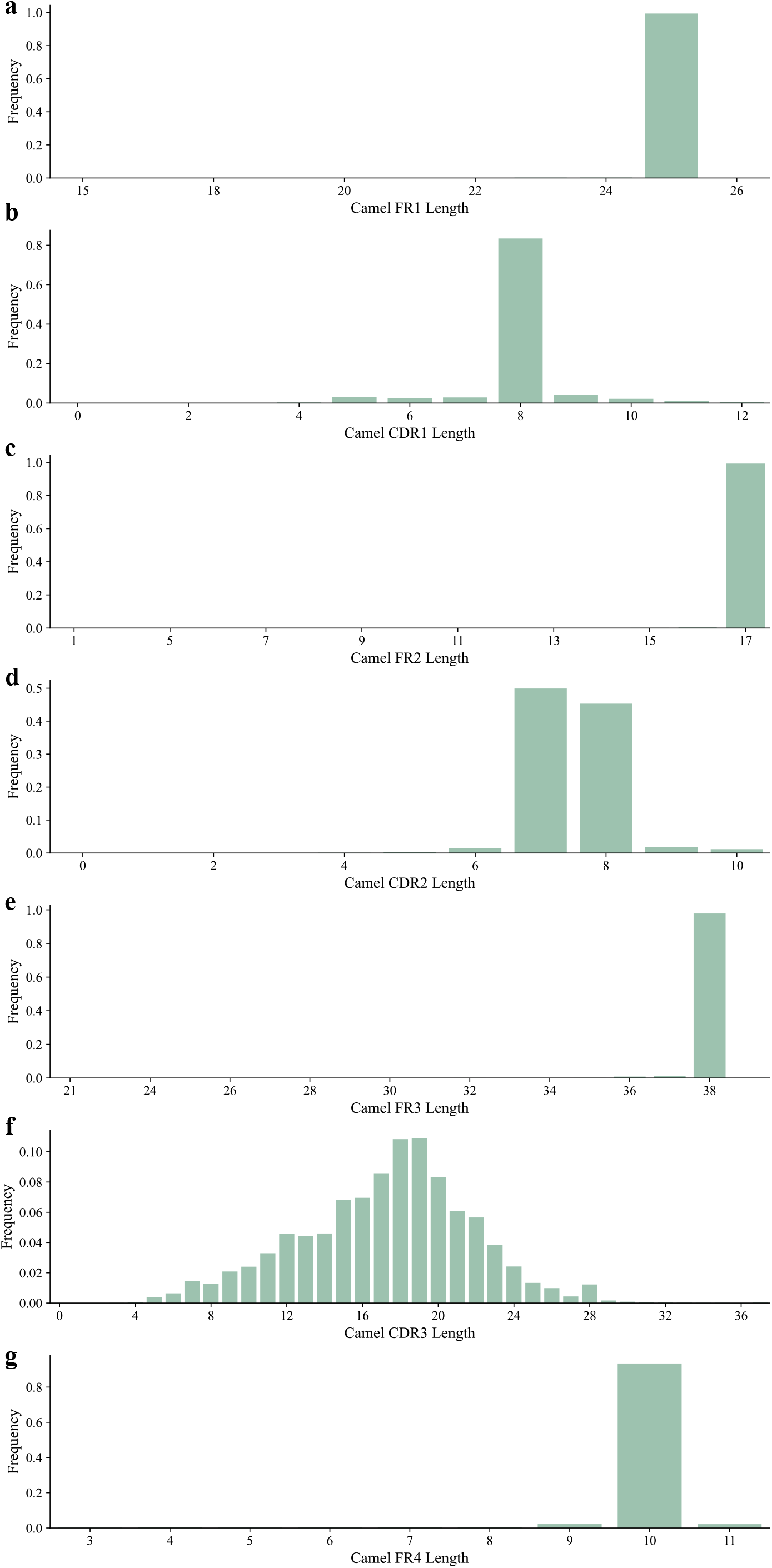
Length distribution of different regions in VHH sequences. The seven frequency histograms represent the length distribution for FR1, CDR1, FR2, CDR2, FR3, CDR3, and FR4, respectively, from top to bottom. All length frequencies are derived from IMGT-aligned VHH sequences.

**Figure S4.**
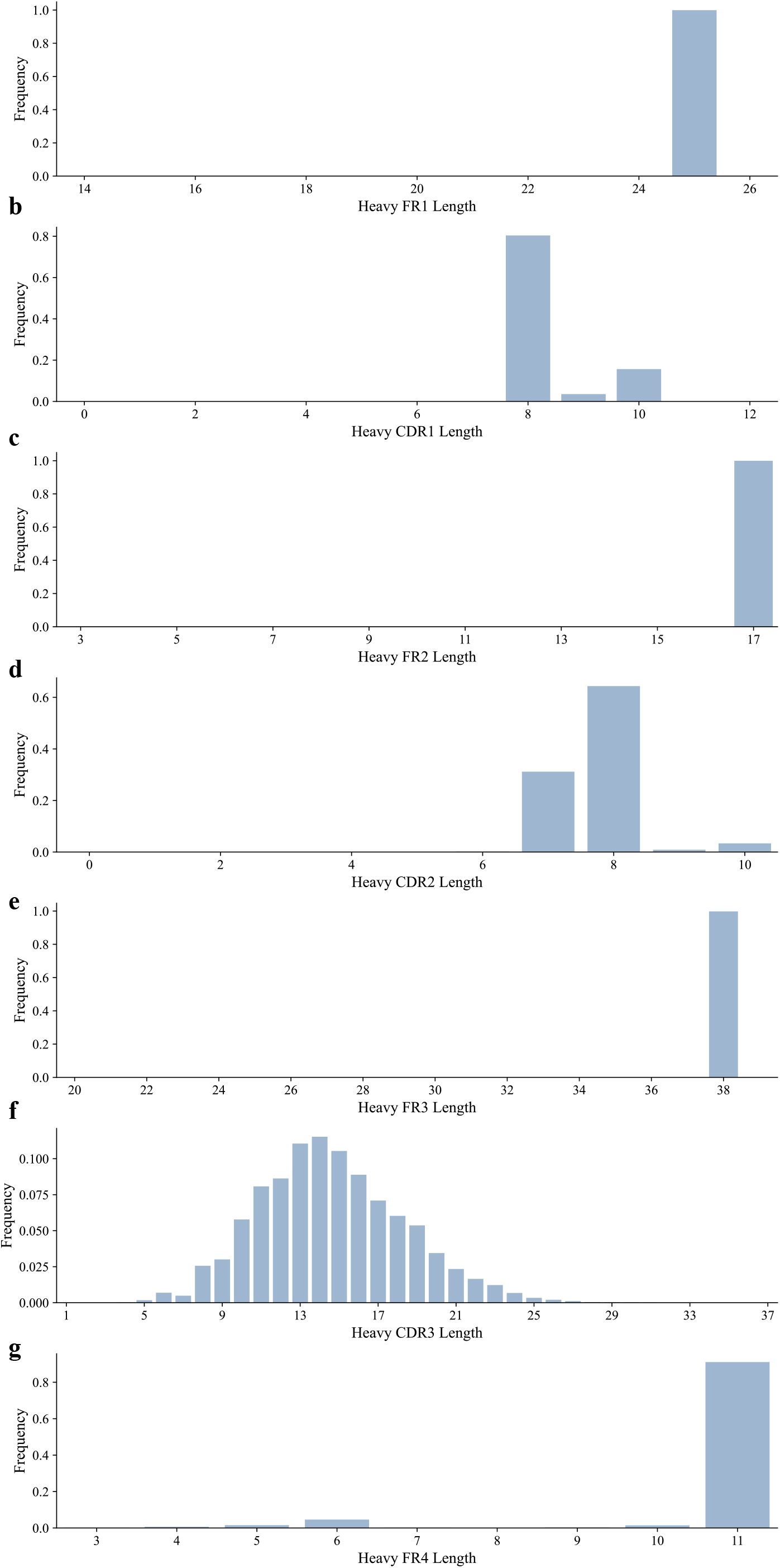
Length distribution of different regions in VH sequences. The seven frequency histograms represent the length distribution for FR1, CDR1, FR2, CDR2, FR3, CDR3, and FR4, respectively, from top to bottom. All length frequencies are derived from IMGT-aligned VH sequences.

**Figure S5.**
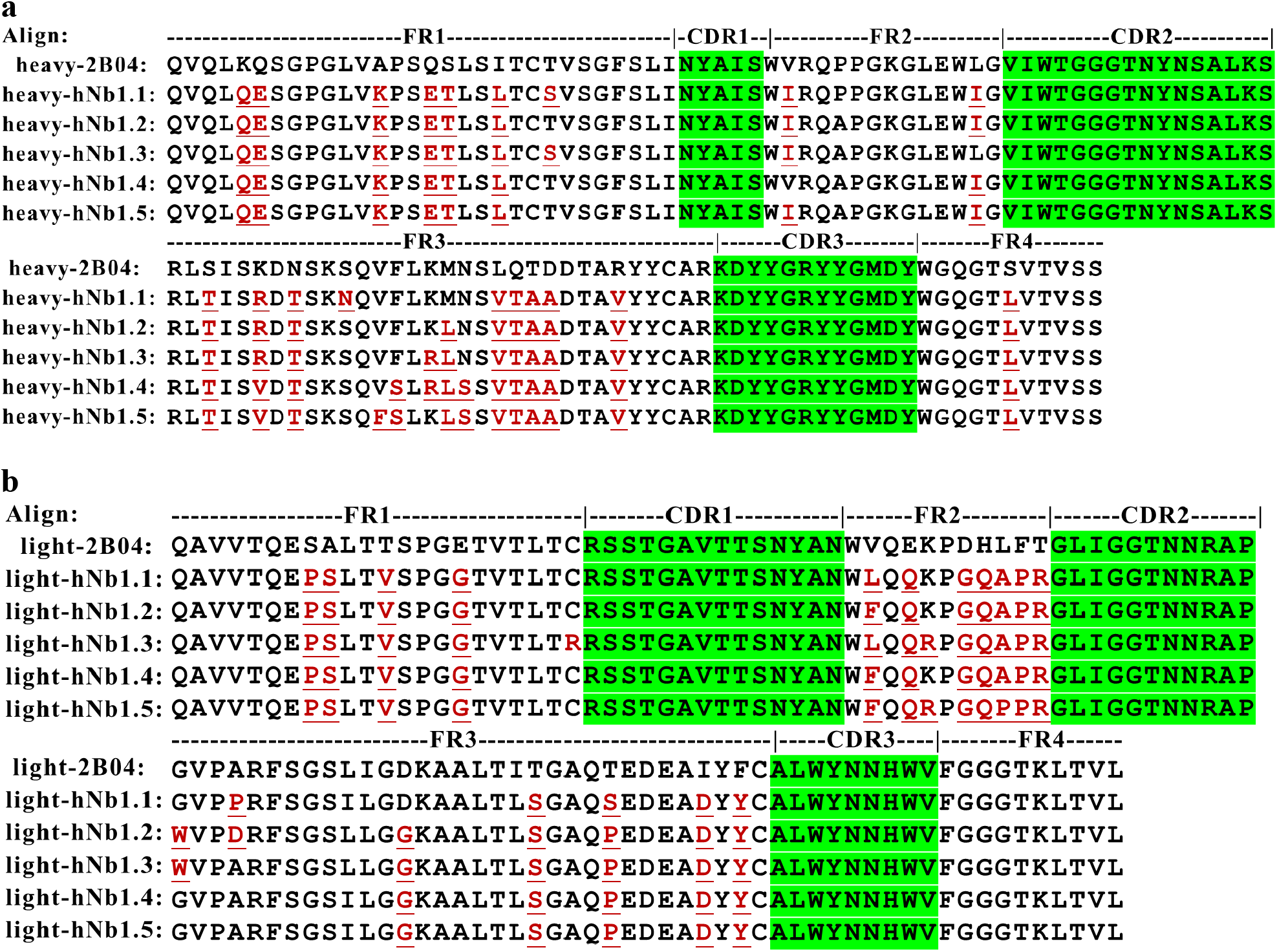
Sequence alignment of humanized antibodies. **a**, Alignment of the heavy chains of humanized antibodies, with residues in the CDR regions highlighted in green. The underlined residues in red indicate the mutations introduced in the humanized heavy chain compared to the parental antibody’s heavy chain. **b**, Alignment of the light chains of humanized antibodies, with residues in the CDR regions highlighted in green. The underlined residues in red denote the mutations introduced in the humanized light chain compared to the parental antibody’s light chain.

**Figure S6.**
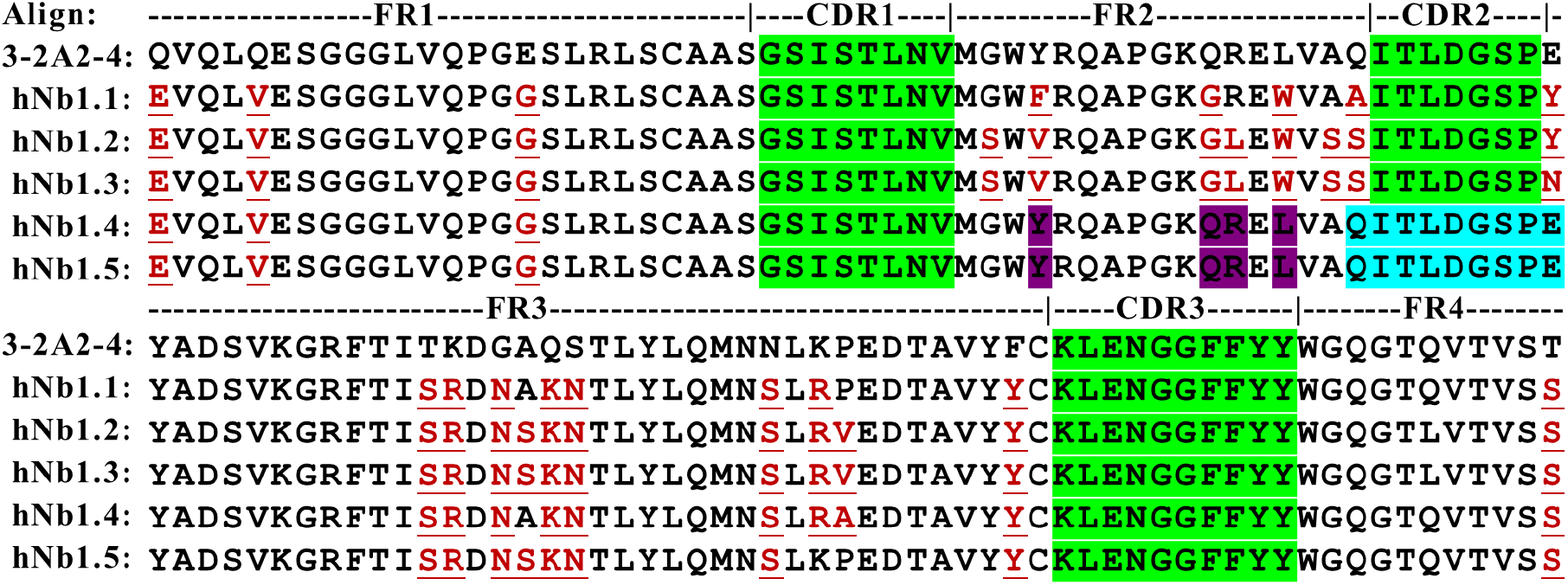
Sequence alignment of humanized nanobodies. In the sequence alignment, residues in the CDR regions are highlighted in green. Light blue indicates the residues defined as CDR2, following the definition used by Llamanade. Key residues preserved by the ‘inp’ sampling method are shown in purple. The underlined residues in red represent the mutations introduced in the humanized nanobodies compared to the parental nanobody.

**Figure S7.**
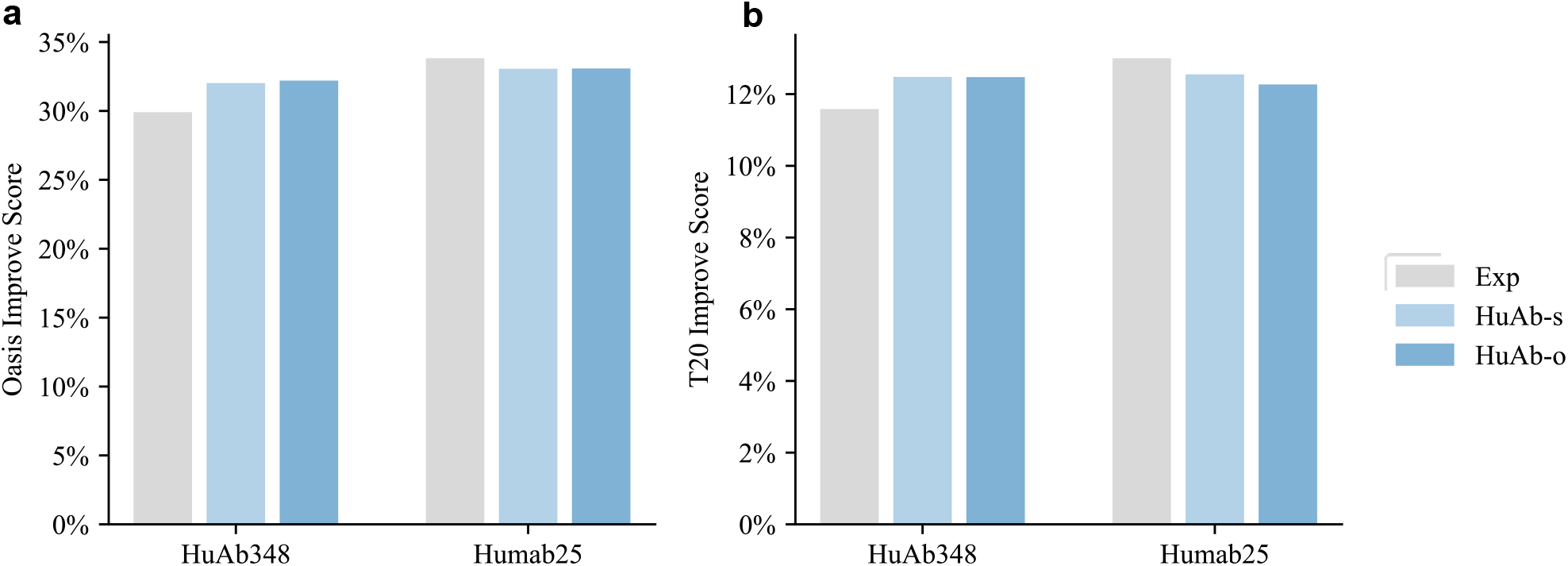
Comparison of sampling order. **a**, OASis score performance for the shuffled sampling order of HuDiff-Ab (denoted as HuAb-s) and the left-to-right sampling order of HuDiff-Ab (denoted as HuAb-o) on the HuAb348 and Humab25 datasets. The grey bars represent the OASis score performance of experimentally humanized sequences for each dataset. **b**, T20 score performance for the shuffled sampling order of HuDiff-Ab (denoted as HuAb-s) and the left-to-right sampling order of HuDiff-Ab (denoted as HuAb-o) on the HuAb348 and Humab25 datasets. The grey bars represent the T20 score performance of experimentally humanized sequences for each dataset.

**Figure S8.**
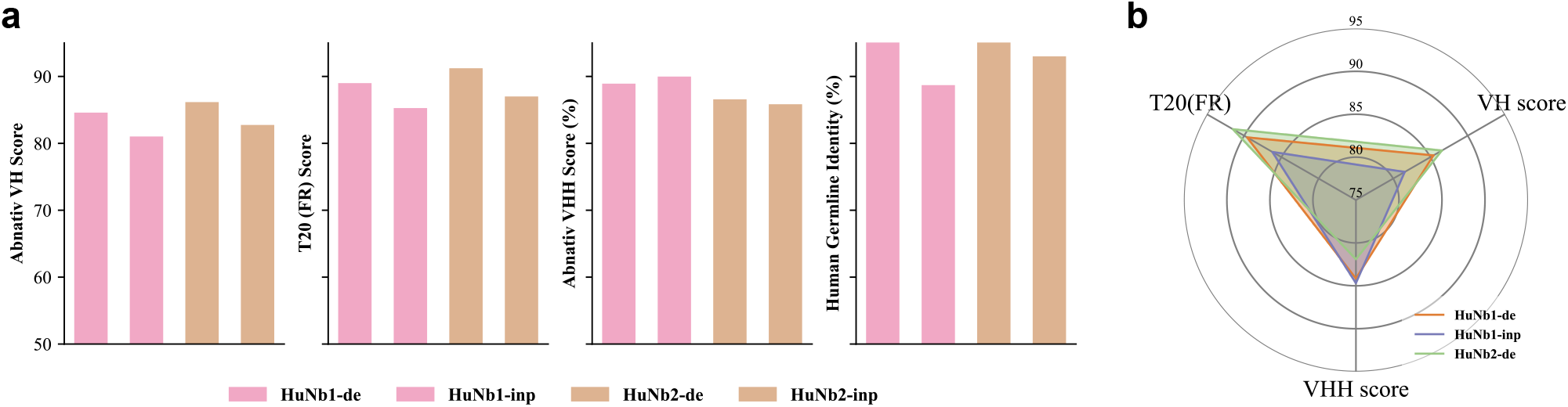
Ablation study of HuDiff-Nb. **a**, Comparison of the AbNativ VH score, T20 (FR), AbNativ VHH score, and germline identity across different versions of HuDiff-Nb model on the Nano300 dataset. HuNb1 represents the HuDiff-Nb model fine-tuned using both the VH and VHH scores from the AbNativ models, while HuNb2 represents the HuDiff-Nb model fine-tuned using only the VH score. **b**, Performance of HuNb1-de, HuNb2-de, and HuNb2-inp on the Nanobert18 dataset, indicating that HuNb1-de consistently outperforms other two versions in T20 and VH scores.

**Figure S9.**
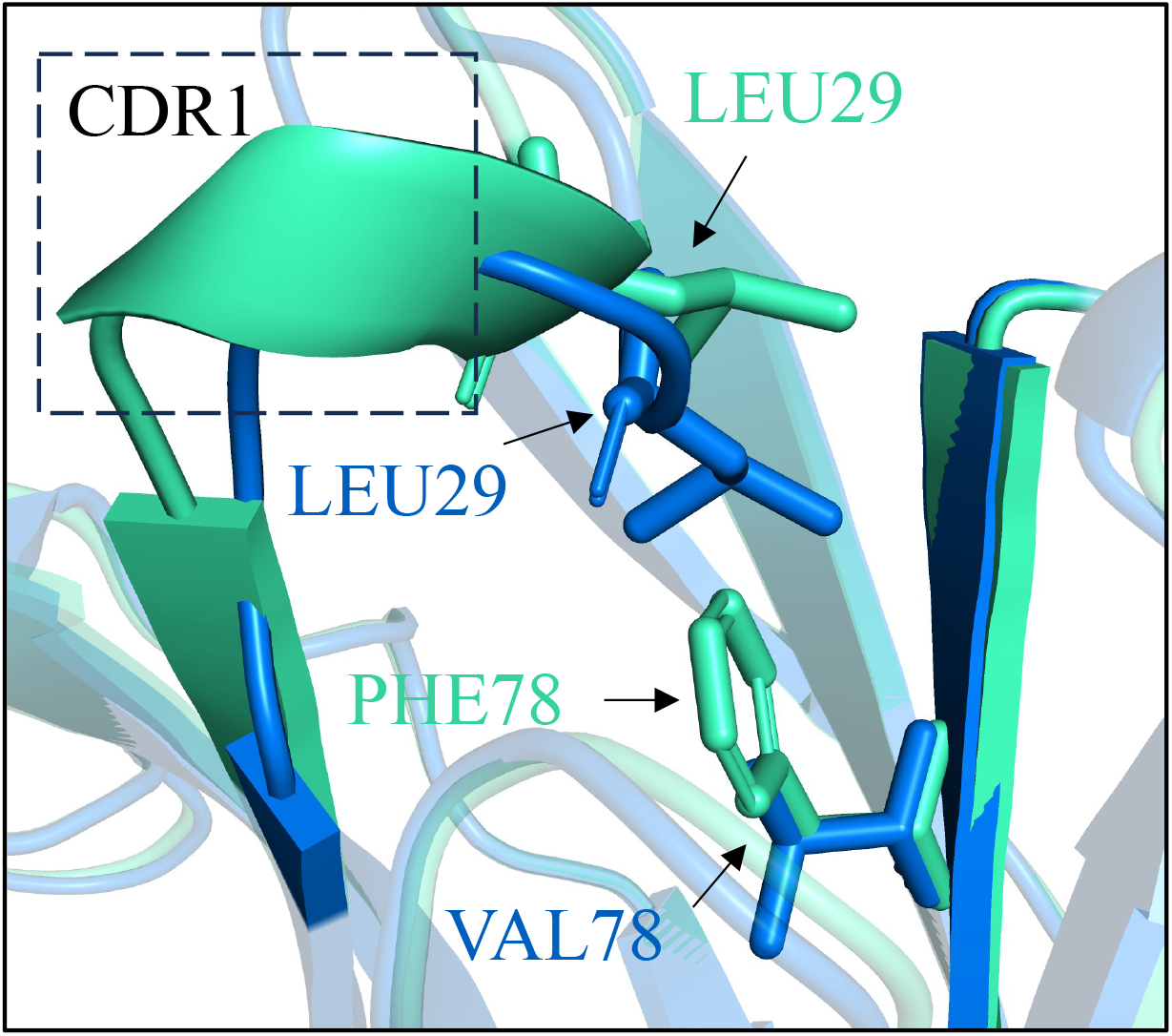
Structural exploration of discrepancy in binding affinity. The blue cartoon structure represents hAb1.4, which has a binding affinity of 1.03 nM, while the green cartoon structure represents hAb1.5, with a binding affinity of 6.37 nM. In hAb1.4, no steric hindrance is observed between residues VAL78 and LEU29. However, in hAb1.5, the bulkier phenyl ring of residue PHE78 creates steric repulsion with LEU29, leading to a conformational change in CDR1.

**Table S1.**
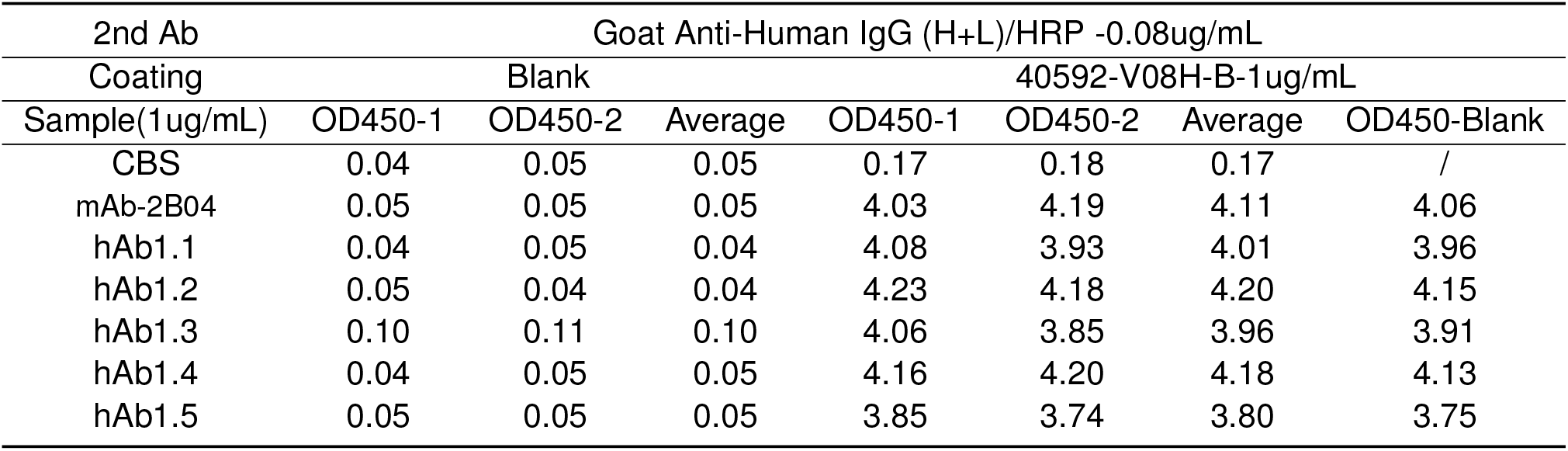
ELISA experiment showing OD450 values for various antibody samples. The table includes readings for blank controls and samples coated with 1 μg/mL of each antibody, measured with a secondary antibody (Anti-Human IgG (H+L)/HRP at 0.08 ug/mL) and the target antigen (1 μg/mL). OD450 values are provided for two replicates (OD450-1 and OD450-2) and their averages.

**Table S2.**
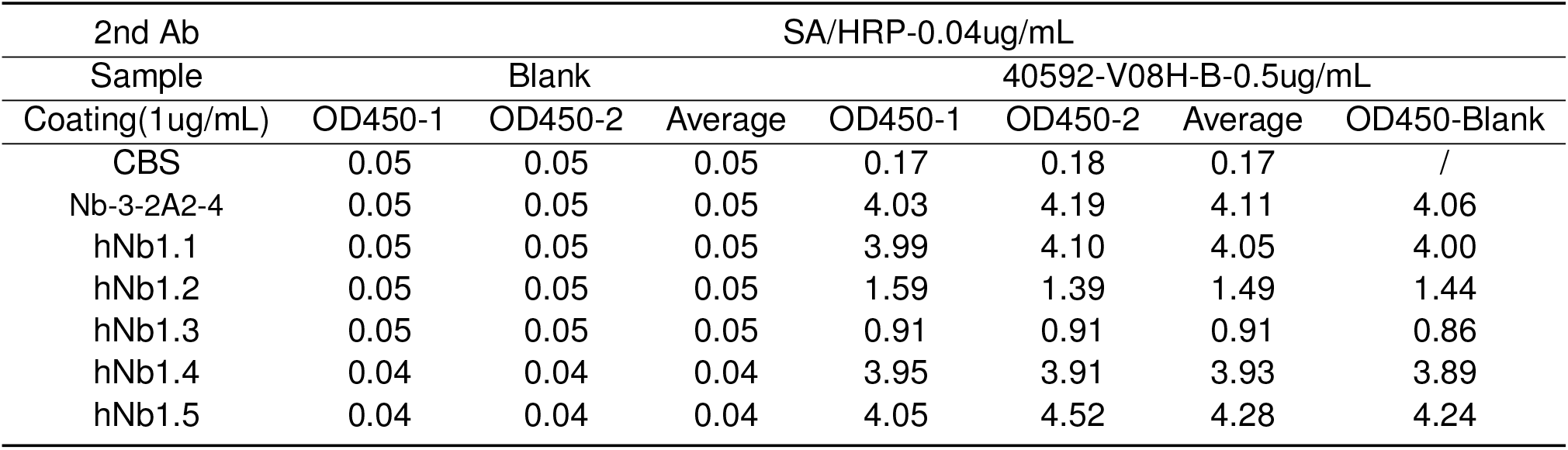
ELISA experiment showing OD450 values for various nanobody samples. The table includes readings for blank controls and samples coated with 1 μg/mL of each nanobody, measured with a secondary antibody (SA/HRP at 0.04 μg/mL) and the target antigen (0.5 μg/mL). OD450 values are provided for two replicates (OD450-1 and OD450-2) and their averages.

**Table S3.** Performance comparison among humanized sequences. This table compares the performance of humanized sequences generated by our method, experimental humanized sequences, and Sapiens1 humanized sequences on the Putative dataset, focusing on improvements in OASis and T20 scores, as well as the germline identity of the framework region (FR). The bold text highlights the highest value for each corresponding metric.

| Data | Method | Humanness improvement |  | Germline identity |  |
| --- | --- | --- | --- | --- | --- |
|  |  | Oasis | T20 | FR |  |
|  |  |  |  | H | L |
| Putative Data | Mouse | - | - | 73.74% | 78.04% |
|  | Experiment | 29.62% | **11.04%** | 88.99% | **92.70%** |
|  | HuDiff-Ab | **30.60%** | 10.36% | **91.65%** | 87.22% |
|  | Sapiens1 | 29.47% | 9.88% | 87.12% | 89.58% |

## Footnotes

1 https://absd.pasteur.cloud

2 https://github.com/Merck/BioPhi

3 https://github.com/sangzhe/Llamanade

